# FGF signaling dynamics regulates epithelial patterning and morphogenesis

**DOI:** 10.1101/2020.11.17.386607

**Authors:** Jakub Sumbal, Tereza Vranova, Zuzana Koledova

## Abstract

Single cell assays revealed that growth factor signaling dynamics is actively sensed by a cell and ultimately controls cell fate. However, the effects of growth factor signaling dynamics at the tissue level have been unknown. We used mammary epithelial organoids, time-lapse imaging, fibroblast growth factor 2 (FGF2) variants of different stabilities, mathematical modeling, and perturbation analysis to study the role of FGF2 signaling dynamics in epithelial morphogenesis. We found that fluctuant and sustained FGF signaling dynamics induced distinct morphological and functional states of mammary epithelium through differential employment of intracellular effectors ERK and AKT. ERK activity domains determined epithelial branch size, while AKT activity drove epithelial stratification. Furthermore, FGF signaling dynamics affected epithelial tissue mechanoresponsiveness to extracellular matrix, thereby impinging upon branch elongation. Our study provides new insights into regulation of epithelial patterning and branching morphogenesis by FGF signaling dynamics and into downstream signaling effectors that regulate cellular outcomes.

## Introduction

Orchestration of complex cell behaviors, such as proliferation, migration, differentiation, and death on population level is essential for building functional tissues during morphogenesis. It is achieved through cell-to-cell communication using core signaling pathways, including receptor tyrosine kinase (RTK) signaling, transforming growth factor signaling, WNT, NOTCH and Hedgehog signaling. Spatial and temporal distribution of various ligands and receptors forms the basic molecular infrastructure for signaling. However, it has long been elusive how activation of a specific receptor translates into a ligand-specific response using just a handful of downstream signaling modules that are common for all receptors within particular signaling pathway family, such as ERK, AKT, STAT and PLCγ in RTK signaling pathways, and how these signaling modules are employed and coordinated in multicellular tissues during formation of complex tissue shapes.

Seminal studies in PC12 cell line revealed that on cellular level, different growth factors acting through different RTKs encode different cellular outcomes by inducing distinct temporal patterns of ERK activity - transient or sustained (Gotoh et al., 1990; Nguyen et al., 1993; Santos et al., 2007). Further works elaborated these observations and found that ERK activity dynamics, defined by duration, magnitude, time-course and spatial localization of ERK activity (Muta et al., 2019), acts as a signaling code that defines cell fates (Blum et al., 2019; Ryu et al., 2015). The patterns of ERK activity dynamics are interpreted into cellular outcomes using hierarchical control of gene expression by ERK-regulated transcription factors (Gille et al., 1995; Murphy et al., 2002; Nakakuki et al., 2010). Moreover, the variety of cellular outcomes in response to different growth factors is further increased by differential employment of other downstream signaling pathways, such as AKT, which modulate activity of the effector molecules (Sampattavanich et al., 2018). Current evidence suggests that the sum effect of ligand identity, concentration, temporal dynamics, and combinations with other ligands is sensed, processed and interpreted in a cell-type and context dependent manner(Li and Elowitz, 2019). However, it remains poorly understood how these signaling variables are interpreted and how these cellular outcomes are coordinated on a tissue level during morphogenesis.

Fibroblast growth factor (FGF) signaling is a crucial pathway that regulates vertebrate development from the earliest embryonic stages throughout lifetime. Regulating cell proliferation, survival, differentiation and migration, it controls a wide range of biological functions (Turner and Grose, 2010). Importantly, FGF signaling has a conserved role in regulation of branching morphogenesis from Drosophila to vertebrates, governing development of branched organs such as fly trachea or mammalian lung, kidney, and mammary glands (Lu and Werb, 2008). In mammals, canonical FGF signaling pathway comprises of 15 extracellularly secreted FGFs, which bind to seven isoforms of FGF receptors (FGFRs). Binding of FGF ligand to FGFR results in receptor dimerization, phosphorylation and activation of downstream signalling pathways, including ERK, AKT, PLCγ and STAT3 signaling pathways (Turner and Grose, 2010).

Changes in FGF ligand expression, retention, and diffusion, including formation of gradients, and fluctuations in FGF signaling are inherent to mammalian development (Balasubramanian and Zhang, 2016; Makarenkova et al., 2009; Niwa et al., 2007; Ornitz and Itoh, 2015; Wahl et al., 2007). However, it remains incompletely understood how FGF signaling dynamics regulates morphogenesis. Therefore, in this work, we used a well-established experimental model of FGF2-induced branching morphogenesis of primary mammary epithelial organoids (Ewald et al., 2008) to study the role of FGF signaling dynamics in epithelial morphogenesis. To induce various FGF ligand dynamics, we used two variants of FGF2, the wild-type FGF2 (FGF2-wt) and its hyperstable mutant (FGF2-STAB) created by protein engineering (Dvorak et al., 2018). While FGF2-wt is naturally unstable and undergoes fast thermal unfolding and deactivation (half-life approximately 6 h at 37°C), FGF2-STAB exhibits high thermal stability and activity over 30 days at 37°C (Dvorak et al., 2018; Koledova et al., 2019). We applied these ligands in a range of concentrations and medium changing strategies, which created a variety of FGF ligand availability schemes according to mathematical modelling. Using these methodological approaches, we provide new insights into regulation of epithelial patterning and branching morphogenesis by FGF signaling dynamics and the specific roles of downstream signaling components in regulation of distinct morphological outcomes.

## Results

### FGF signaling dynamics governs epithelial morphogenesis

To assess the role of FGF signaling dynamics in epithelial morphogenesis, we treated mammary organoids with 1 nM FGF2-wt, which we delivered in a range of different medium changing strategies over 9 days of culture (Supplemental Figure 1A). The most commonly used strategy in cell culture, with the medium change every three days (C3d), induced formation of thin branches (i.e. normal branching of organoids) as expected (Figure 1A, B). When the fresh FGF2-wt was added only once at the beginning of culture and was left for the duration of culture with no change (NC) of the medium, or was added only for one- or three-hour pulse (p1h, p3h), no branching was observed, and organoids only mildly grew and formed cysts (Figure 1A). Increasing frequency of medium change to every day (Ced) or every 6 hours (C6h) increased the percentage of branching organoids (Figure 1A, B) and the phenotype of branching was the same as when the medium was changed every three days. But when we added fresh FGF2-wt every 6 hours (A6h), which is the half-life of FGF2-wt, a new epithelial phenotype emerged, characterized by thick branches, that we named massive branching (Figure 1A, B). Mathematical modeling of FGF2 concentration in the medium revealed that the condition of adding 1 nM FGF2 every 6 hours was the only condition when the concentration of FGF2-wt in the medium did not drop much below 1 nM during the whole organoid culture period (Figure 1C). This suggested that sustained signaling of 1 nM FGF2 is critical for formation of massive branches.

**Figure 1.**
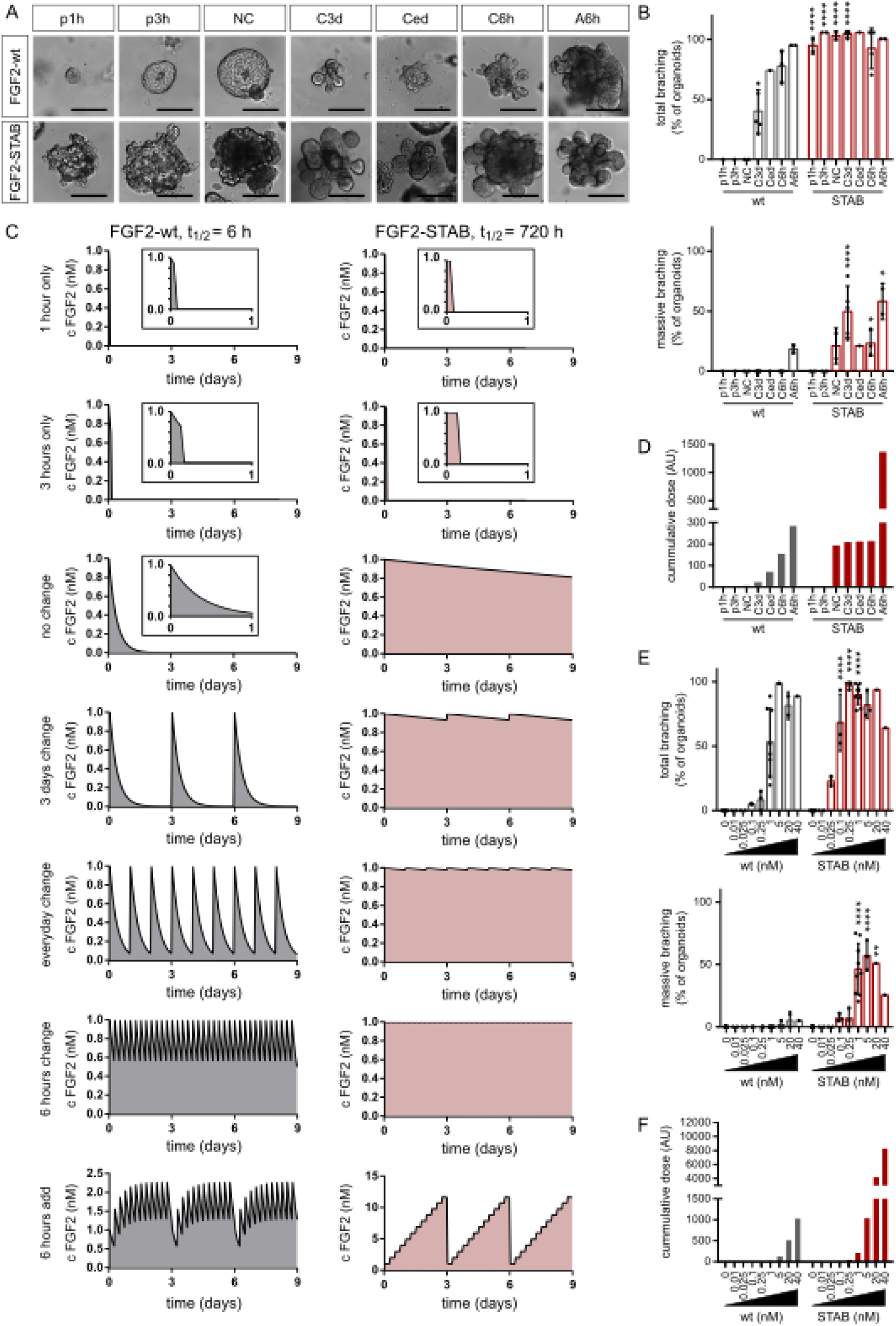
FGF signaling dynamics governs epithelial morphogenesis. **A**. Organoids after 9 days of culture with different FGF signaling dynamics as depicted in **C**. Scale bars, 200 μm. P1h, 1-hour pulse; p3h, 3-hour pulse; NC, medium was not changed during the whole 9-day culture of organoids; C3d, medium was changed every 3 days; Ced, medium was changed every day; C6h, medium was changed every 6 h; A6h, fresh FGF2 was added to the medium every 6 h and every 3 days, the full medium volume was changed. **B**. Total (normal + massive) branching and massive branching of organoids subjected to different medium changing strategies. The plots show mean + s.d., N = 20-100 organoids, n = 1-5 independent experiments; for exact numbers see Supplemental Table 4. The asterisks indicate significant difference between FGF2-wt and FGF2-STAB per condition; *P < 0.05; ****P < 0.0001 (two-way ANOVA). **C**. The graphs show mathematic models of FGF2 concentration dynamics in culture during different medium changing strategies. The models are based on the FGF2 half-lives (6 h for FGF2-wt and 720 h for FGF2-STAB). Potential active FGF2 degradation or production by the cells was not accounted for. FGF2-wt is depicted in grey, FGF2-STAB in pink. Insets show detail of FGF dynamic during the first day of culture. **D**. The plot shows cumulative FGF2 dose, calculated from the mathematic model of FGF2 concentration dynamics as the area under the curves shown in **C**. **E**. Analysis of organoid morphogenetic response to a range of FGF2 concentrations. The plots show quantification of total (normal + massive) branching and massive branching of organoids, respectively, in response to a range of FGF2 concentrations. The plots show mean + s.d., N = 20-180 organoids, n = 1-9 independent experiments; for exact numbers see Supplemental Table 4. The asterisks indicate significant difference between FGF2-wt and FGF2-STAB per condition; **P < 0.01; ****P < 0.0001 (two-way ANOVA). **F**. The plot shows cumulative FGF2 dose, calculated from the mathematic model of FGF2 concentration dynamics as the area under the curve.

To test this finding, we employed a stabilized form of FGF2 with long-term thermostability, FGF2-STAB (Dvorak et al., 2018). Unlike FGF2-wt, FGF2-STAB remains active after preincubation at 37°C for 7 days and effectively induces branching in the mammary organoids system (Supplemental Figure 2A, B). We applied the same medium-changing or FGF2-adding strategies with FGF2-STAB as we had done with FGF2-wt. In all these conditions, FGF2-STAB effectively induced epithelial branching, and in most of the conditions FGF2-STAB induced the massive branching phenotype (Figure 1A-C). Only when the FGF2-STAB was applied in a one- or three-hour pulse, the organoids developed thin branches (Figure 1A, B). Importantly, these were the only two conditions in which the FGF2 concentration dropped significantly below 1 nM over the organoid culture period (Figure 1C).

### Mathematic modeling reveals a critical dose of FGF signaling for induction of massive branches

To express mathematically the amount of FGF signaling achieved by FGF2 supplied at a known concentration by different medium-changing or FGF2-adding strategies over the organoid culture period, we calculated a “cumulative dose” for each condition. The cumulative dose corresponds to the area under the curve of the plot modeling FGF2 dynamics over time (see Methods). These calculations revealed that for FGF2-wt the cumulative dose of FGF signaling increased with increasing frequency of medium change or FGF2 addition. In the case of FGF2-STAB, the cumulative dose of FGF signaling was relatively stable from the no medium change to the medium change every 6 hours strategy (Figure 1D). Adding FGF2-STAB every 6 hours led to a major increase of the cumulative dose but it had no further effect on the epithelial phenotype, suggesting that the system was already saturated. Importantly, these calculations further showed that only the condition of adding FGF2-wt every 6 hours, which was the only one inducing massive branching with FGF2-wt, reached the cumulative dose of FGF signaling similar to the dose achieved by FGF2-STAB at conditions when it induced massive branching (Figure 1D).

Furthermore, we exposed the organoids to a wide range of FGF2-wt or FGF2-STAB concentrations under the strategy of medium change every three days. With the increasing concentration of FGF2 the percentage of branching organoids increased for both FGF2-wt and FGF2-STAB, and FGF2-STAB showed a ten times higher total branching inducing potency than FGF2-wt (Figure 1E). FGF2-STAB very effectively induced massive branching of organoids from 1 nM concentration. FGF2-wt induced massive branching only at 20 nM or higher concentrations, yet still only in a low percentage of organoids. The efficiency of induction of massive branching correlated with the cumulative dose of FGF signaling (Figure 1F).

Taken together, we found that not only the FGF2 concentration per se, but the temporal dynamics of FGF2 signaling regulates distinct morphogenetic outcomes in the mammary epithelium. When 1 nM FGF2 is supplied, fluctuant FGF signaling induces normal branching, while sustained FGF signaling induces massive branching, as a result of the cumulative dose of FGF acquired over time.

### FGF signaling dynamics regulates multiple epithelial cell functions that contribute to the morphogenetic outcome

Next, we sought to characterize the new massive branching phenotype, induced by sustained FGF signaling. To this end, we cultured organoids with 1 nM FGF2-wt or FGF2-STAB with medium changed every three days. By careful analysis of time-lapse movies of growing organoids, we found that in massively branching organoids, the branching occurs later than in normally branching organoids and is preceded by an enormous growth of the epithelium (Figure 2A, B; Supplemental Figure 3A, Supplemental Videos 1-3). Histological analysis of the organoids showed that the massive branches induced by sustained FGF signaling were formed by prominently stratified epithelium (Figure 2C; Supplemental Figure 3B). Moreover, the normally and massively branched organoids differed in epithelial cell type distribution. The normal branches of organoids exposed to fluctuant FGF signaling lacked myoepithelial cells at the tips of the branches, a phenomenon previously reported for mammary organoids cultured in Matrigel (Nguyen-Ngoc and Ewald, 2013). However, the massive branches of FGF2-STAB-treated organoids were fully covered by myoepithelial cells (Figure 2C, D; Supplemental Figure 4A, B).

**Figure 2.**
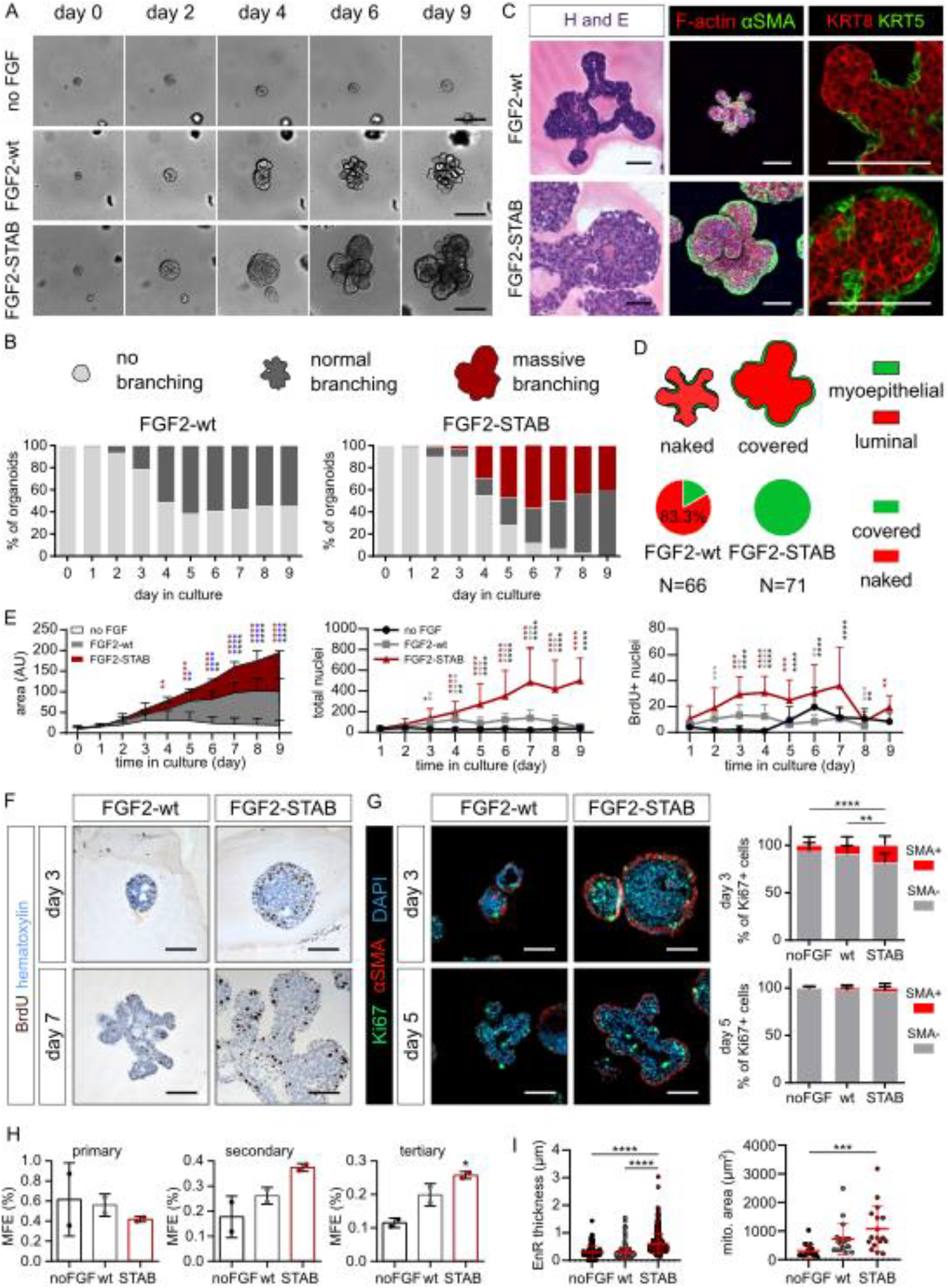
Sustained FGF signaling regulates multiple epithelial cell functions. **A**. Morphogenesis of organoids with no FGF, 1 nM FGF2-wt, or 1 nM FGF2-STAB. Snapshots are from a time-lapse experiment (see also Supplemental Videos 1-3). Scale bars, 200 μm. **B**. Temporal analysis of organoid morphogenetic phenotypes. Light gray, no branching; dark gray normal branching; dark red, massive branching. A sum of three independent experiments with 20 organoids tracked per condition per experiment. **C**. Whole-mount, histology and immunofluorescence of FGF2-wt- or FGF2-STAB-treated organoids. αSMA, α smooth muscle actin; H and E, hematoxylin and eosin; KRT5, keratin 5; KRT8, keratin 8. Scale bars, 100 μm. **D**. Quantification of branch coverage with αSMA+ cells. **E**. Quantification of organoid growth from time-lapse experiments and from total number of nuclei or BrdU+ nuclei (shown in **B** and Supplemental Figure 5A). The plot shows mean + s.d., n = 3, N = 60 organoids (time-lapse) and N = 3-40 organoids (histology) per condition; for exact numbers see Supplemental Table 5. Red asterisks indicate the difference between FGF2-STAB and no FGF, blue asterisks between FGF2-wt and no FGF, black asterisks between FGF2-wt and FGF2-STAB. *P < 0.05; **P < 0.01; ***P < 0.001; ****P < 0.0001 (two-way ANOVA for time-lapse experiment; multiple t-tests with Holm-Sidak’s test for histology). **F**. BrdU staining of organoids treated with FGF2-wt or FGF2-STAB for 3 or 7 days. Scale bars, 50 μm. **G**. Ki67 and αSMA staining of organoids treated with FGF2-wt or FGF2-STAB for 3 or 5 days and quantification of αSMA-/ αSMA+ cell proportion among Ki67+ cells. Scale bars, 50 μm. The plots show mean + s.d., N = 11-50 organoids per condition (see Supplemental Table 5). ** P < 0.01, ****P < 0.0001 (two-way ANOVA). **H**. Endoplasmic reticulum (EnR) thickness and mitochondrial area of organoids. The plots show mean ± s.d. N = 116, 234, and 436 of ER cisternae, and 17, 19, and 17 mitochondria from organoids treated with no FGF, FGF2-wt, or FGF2-STAB for 5 days, respectively. ***P < 0.001; ****P < 0.0001 (one-way ANOVA). **I**. Primary, secondary and tertiary mammosphere formation efficiency (MFE) of cells from organoids cultured with no FGF, FGF2-wt, or FGF2-STAB for 5 days. The plots show mean + s.d., n = 2 independent experiments. *P < 0.05 (one-way ANOVA).

Besides altering the shape of organoids, sustained FGF signaling also promoted formation of bigger organoids (Figure 2E). Because in the bright field images the real difference in organoid growth could be masked by lumen enlargement, we also counted the total number of nuclei and the number of BrdU+ nuclei per organoid section to measure organoid proliferation, and we assessed apoptosis using staining for cleaved caspase 3. We found significantly increased number of total nuclei as well as BrdU+ nuclei in FGF2-STAB organoids in comparison to FGF2-wt organoids (Figure 2E, F; Supplemental Figure 5A) and decreased apoptosis in FGF2-STAB organoids (Supplemental Figure 5B, C). The most significant difference in BrdU+ nuclei numbers was found on days 3 and 4, which is when the concentric growth of FGF2-STAB-treated organoids is observed. On day 3 we detected an increased proportion of myoepithelial cells among proliferating cells in FGF2-STAB-treated organoids (Figure 2G; Supplemental Figure 6A), which helps to explain the supply of cells for the full myoepithelial coverage of massively branched organoids (Figure 2C, D).

To assess the effect of different FGF signaling dynamics on cell renewal capacity, we dissociated organoids exposed to either no, fluctuant, or sustained FGF signaling to single cells, which we further tested in mammosphere formation assay (Supplemental Figure 7A) or organoid formation assay (Supplemental Figure 7B). Cells derived from FGF2-STAB-treated organoids formed significantly more mammospheres in the third generation (Figure 2H), indicating a higher content of stem cells in FGF2-STAB-treated organoids. In the long-term organoid formation experiment, FGF2-STAB induced formation of bigger and branched organoids, while organoids formed with FGF2-wt were smaller and cystic (Supplemental Figure 7B-D). This suggests that sustained FGF signaling is essential to retain morphogenetic capacity of epithelial cells. Organoids subjected to sustained FGF signaling also showed reduced epithelial polarity, as shown by β-catenin and E-cadherin staining and by ultrastructural analysis of cell-cell connections (Supplemental Figure 8A-C). Additionally, ultrastructural analysis revealed enlarged mitochondria and dilated endoplasmic reticulum in FGF2-STAB treated organoids (Figure 2I; Supplemental Figure 8D), indicating increased metabolic activity of cells subjected to sustained FGF signaling.

Together our data show that FGF signaling dynamics regulates a plethora of cell functions, which on tissue level result in different morphogenic outcomes. In comparison to fluctuant FGF signaling, sustained FGF signaling promotes more cell proliferation, including in myoepithelial cells, which contributes to full myoepithelial coverage of massive branches, reduced apoptosis, higher metabolism and increased epithelial cell regenerative and morphogenetic potential. However, while these functions regulate cell number and tissue size, they do not explain how different tissue shapes arise upon different FGF signaling dynamics. We hypothesized a role of downstream signaling pathways, including ERK and AKT, in regulation of tissue architecture.

### FGF signaling dynamics regulates epithelial branching via ERK signaling

Organoid branching and branch elongation were demonstrated to be regulated by ERK signaling in response to FGF signaling (Huebner et al., 2016). We hypothesized that ERK signaling regulates branch thickness depending on FGF signaling dynamics. Therefore, we analyzed the effect of ERK signaling inhibition by U0126 on organoid branching morphogenesis upon fluctuant and sustained FGF signaling. U0126 abrogated branching of organoids treated either with FGF2-wt or FGF2-STAB (Figure 3A, B). Interestingly, epithelial stratification and full myoepithelial coverage in response to sustained FGF signaling was not affected by U0126 (Figure 3C; Supplemental Figure 9A). A higher concentration of U0126 was needed to inhibit branching in FGF2-STAB-treated organoids in comparison to FGF2-wt-treated organoids, suggesting higher ERK signaling activity in FGF2-STAB-treated organoids. This was further corroborated by Western blot detection of higher amount of active ERK (phosphorylated ERK, pERK) (Figure 3D) and higher expression of FGF-ERK signaling genes (*Dusp6*, *Etv4* and *Etv5*) by qPCR in FGF2-STAB organoids (Figure 3E).

**Figure 3.**
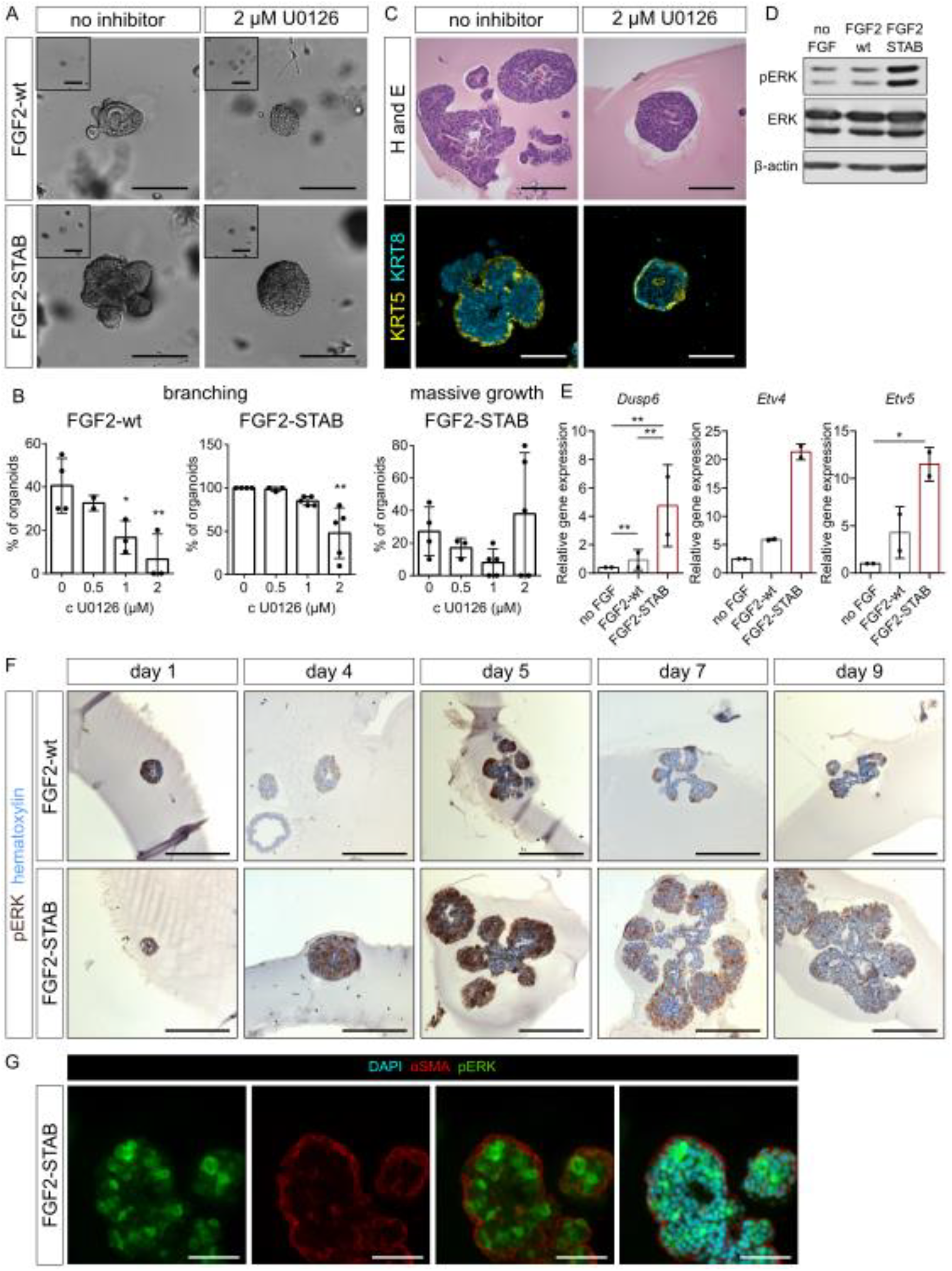
FGF2 signaling dynamics regulates epithelial branching via ERK signaling. **A**. Organoids cultured with FGF2-wt or FGF2-STAB and with or without ERK inhibitor U0126 for 9 days. Insets show organoids on day 0. Scale bars, 200 μm. **B**. Quantification of organoid morphogenetic response – total branching or massive growth – to ERK inhibitor at a range of concentrations. The plots show mean + s.d., n = 2-4 independent experiments, N = 20 organoids per experiment; for exact numbers see Supplemental Table 4. *P < 0.05; **P < 0.01; ***P < 0.001 (one-way ANOVA). **C**. Histological and immunofluorescence analysis of FGF2-STAB-treated organoid architecture upon ERK inihibitor treatment. H and E, hematoxylin and eosin; KRT5, keratin 5; KRT8, keratin 8. Scale bars, 100 μm. **D**. Phosphorylated ERK (pERK) distribution in sections of organoids treated with 1 nM FGF2-wt or FGF2-STAB. Scale bars, 200 μm. **E**. Western blot analysis of pERK, total ERK, and β-actin amount in organoids on day 5 in culture, 48 h after FGF2 treatment. **F**. qPCR analysis of *Dusp6, Etv4* and *Etv5* gene expression in organoids on day 5 in culture, 48 h after FGF2 treatment. The plots show mean + s.d., n = 2. *P < 0.05; **P < 0.01 (one-way ANOVA). **G**. Distribution of pERK in basal (αSMA+) and luminal (αSMA-) cells in a section of organoid treated with FGF2-STAB for 5 days. Scale bars, 50 μm.

Analysis of spatial distribution of pERK in the organoids revealed that in response to sustained as well as fluctuant FGF signaling, pERK is mosaically distributed early during the morphogenesis in round organoids, but forms distinct domains in the tips of the branches, while the necks of the branches are poor in pERK (Figure 3F). In the branches, pERK positive cells are localized mainly in layers of luminal cells (Figure 3G). Importantly, the domains of pERK were bigger in response to sustained FGF signaling, suggesting that the size of pERK domains determines the thickness of the branches (Figure 3F).

### AKT signaling is crucial for epithelial stratification

Besides the ERK pathway, several other signaling pathways act downstream of FGFR, including AKT, STAT and PLCγ pathways. To assess their contribution to the morphogenetic response upon different dynamics of FGF signaling, we used inhibitors of AKT (Akti1/2), STAT3 (Stattic), or PLCγ (U73122). But first as a control that the effects of both FGF2-wt and FGF2-STAB were mediated through FGFR, we used BGJ398, an FGFR inhibitor. BGJ398 effectively inhibited organoid growth and branching morphogenesis in both FGF2-wt- and FGF2-STAB-treated organoids. A higher concentration of BGJ398 was needed to significantly abrogate branching induced by FGF2-STAB than by FGF2-wt, suggesting a higher level of FGFR signaling induced by FGF2-STAB (Supplemental Figure 10A-C). Inhibition of either STAT3 or PLCγ somewhat decreased normal branching by FGF2-wt and massive branching by FGF2-STAB, but without reaching statistical significance (Supplemental Figure 10A-C). However, inhibition of AKT led to a significant decrease in both FGF2-wt- and FGF2-STAB-induced branching (Figure 4A, B) and, importantly, caused a dramatic loss of massive growth and epithelial stratification induced by sustained FGF signaling (Figure 4A-C). Additionally, we detected elevated amount of phosphorylated AKT (pAKT, activated) in FGF2-STAB-treated organoids (Figure 4D), suggesting that sustained FGF signaling leads to hyperactivation of this pathway.

**Figure 4.**
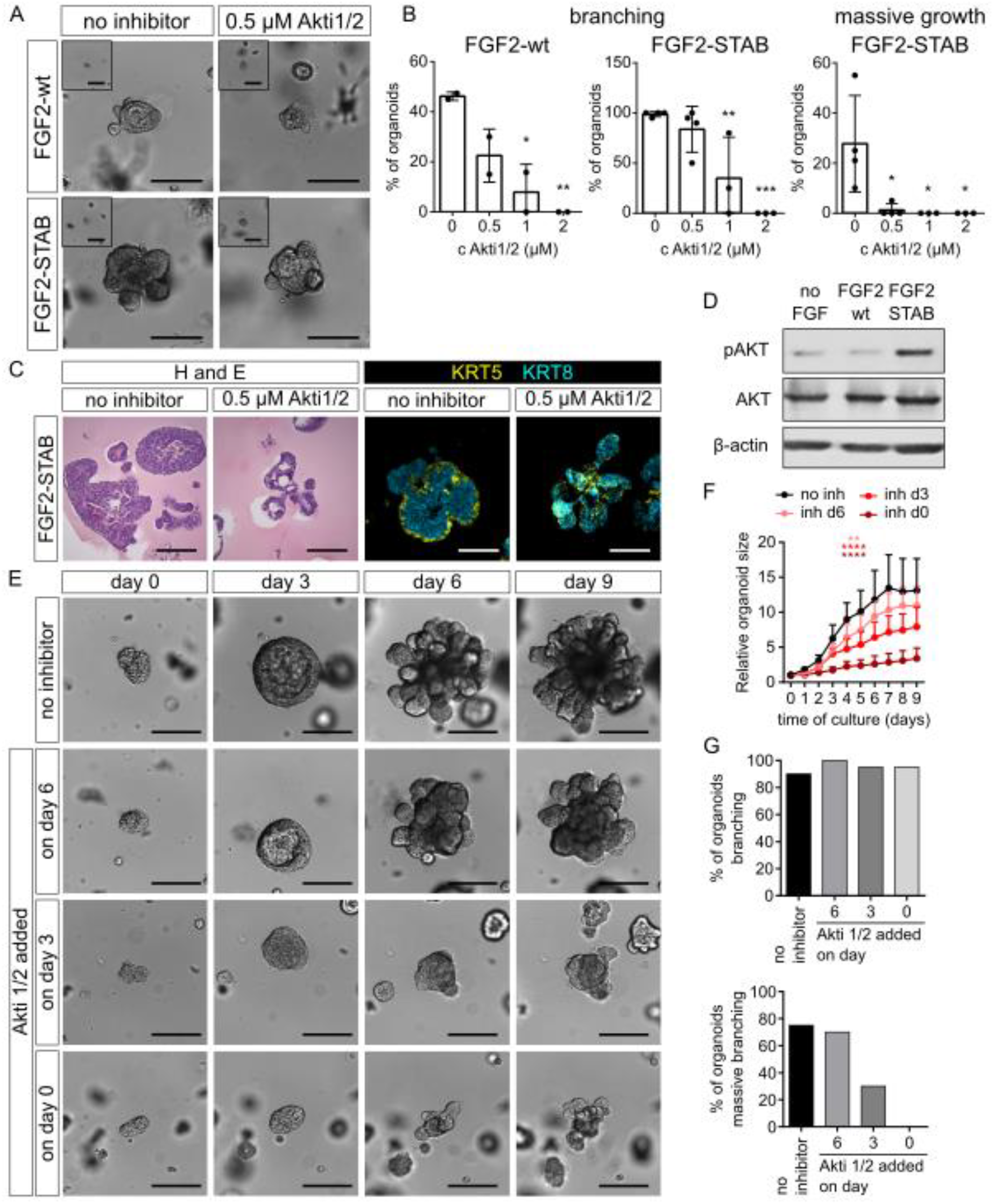
AKT signaling is crucial for sustained FGF2 signaling-induced epithelial stratification. **A**. Organoids cultured with FGF2-wt or FGF2-STAB and with or without AKT inhibitor (Akti) for 9 days of culture. The images of organoids without inhibitor are the same as in Figure 3A because they are from the same experiment. Insets show organoids on day 0. Scale bars, 200 μm. **B**. Quantification of organoid morphogenetic response – total branching or massive growth – to Akti1/2 at a range of concentrations. The plots show mean + s.d., n = 2-5 independent experiments, N = 20 organoids per experiment; for exact numbers see Supplemental Table 4. *P < 0.05; **P < 0.01; ***P < 0.001 (one-way ANOVA). **C**. Histological and immunofluorescence analysis of FGF2-STAB-treated organoid architecture upon Akti treatment. H and E, hematoxylin and eosin; KRT5, keratin 5; KRT8, keratin 8. Scale bars, 100 μm. **D**. Western blot analysis of phosphorylated AKT (pAKT), total AKT and β-actin level in organoids on day 5 of culture, 48 h after treatment with FGF2. **E**. Organoid morphogenesis in response to FGF2-STAB and no inhibitor or AKT inhibitor (0.5 μM Akti1/2) added on day 0, 3 or 6 of culture. The images are snapshots from time-lapse imaging of the organoids. Scale bars, 200 μm. **F**. FGF2-STAB-treated organoid size upon no inhibitor or treatment with Akti1/2 on day 0, 3, or 6. The plot shows mean + s.d., n = 1 experiment, N = 20 organoids per condition. **P < 0.01; ****P < 0.0001 (one-way ANOVA). Pink aterisks: control (no inhibitor) to inhibitor on day 6; red aterisks control to inhibitor on day 3; dark red aterisks control to inhibitor on day 0). **G**. Quantification of organoid total branching and massive growth in cultures from **D**. N = 20 organoids per condition.

Based on our data from time-lapse imaging (Figure 2A, B) we hypothesized that the concentric massive growth of organoids around day 3 is crucial for subsequent massive branching because the biggest differences in new branch development and proliferation occurs at that time. Therefore, to assess whether at that time the AKT signaling is important for the massive growth, we treated organoids under sustained FGF signaling with Akti1/2 from either day 0, 3, or 6. When Akti1/2 was added on day 6, massive branching occurred, similarly to organoids with no inhibitor (Figure 4E, G). When Akti1/2 was added on day 3, the growth of the organoid was severely reduced, similarly to Akti1/2 addition from day 0 (Figure 4E, F). Importantly, although the growth and stratification of the organoid did not occur, development of new branches was not affected (Figure 4E, G). This suggested that AKT signaling is essential for epithelial stratification.

The basal organoid medium, in which the organoids are cultured and exposed to FGF signaling, contains a potent inducer of AKT signaling, insulin. To assess how this additional insulin-induced AKT signaling contributes to organoid morphogenesis, we cultured the organoids under FGF signaling without the insulin-containing component of the basal organoid medium (supplement ITS) or in the presence of insulin receptor inhibitor (BMS 536924). Loss of insulin signaling led to a similar phenotype as the AKT inhibition; total organoid branching was not affected, but the massive growth and epithelial stratification of FGF2-STAB treated organoids were lost completely (Supplemental Figure 11 A-C).

At last we inhibited both AKT and ERK signaling at the same time by combination of Akti1/2 and U0126. This combined inhibition completely abrogated any morphogenesis (Supplemental Figure 12A), similarly to FGFR inhibition (Supplemental Figure 10A-C), suggesting that ERK and AKT pathways are the major pathways orchestrating epithelial morphogenesis downstream of FGF.

### Massive branches induced by sustained FGF2 signaling in ex vivo organoids phenocopy terminal end buds in vivo

During mammary gland development, soluble signals are integrated with mechanical signals to guide morphogenesis (Gjorevski and Nelson, 2011). Therefore, we investigated the effect of different FGF2 signaling dynamics on mammary epithelial morphogenesis in extracellular matrix (ECM) of increased stiffness and fibrillarity – a mixture of Matrigel with collagen I – which is by composition and physical properties closer to ECM in vivo (Nguyen-Ngoc and Ewald, 2013). Concordantly to previous reports (Neumann et al., 2018; Nguyen-Ngoc and Ewald, 2013), we found that in the mixture of Matrigel with collagen, organoids formed significantly longer branches when exposed to fluctuant FGF2 signaling. However, when exposed to sustained FGF2 signaling, the organoid branches did not elongate (Figure 5A, B; Supplemental Videos 4 and 5). This suggested uncoupling of FGF2-STAB-treated organoids from mechanical signals of the ECM. To test if the epithelial cells exert mechanical forces on the surrounding ECM, we cultured the organoids in fluorescently labelled collagen. When cells pull on the collagen fibers, the fibers are aligned closer together, increasing fluorescent signal intensity. Aligned collagen was visible around branch necks of the FGF2-wt-treated organoids, but not around the massively branched FGF2-STAB-treated organoids (Figure 5C), demonstrating lack of mechanotransduction between the FGF2-STAB-treated organoid and the surrounding ECM.

**Figure 5.**
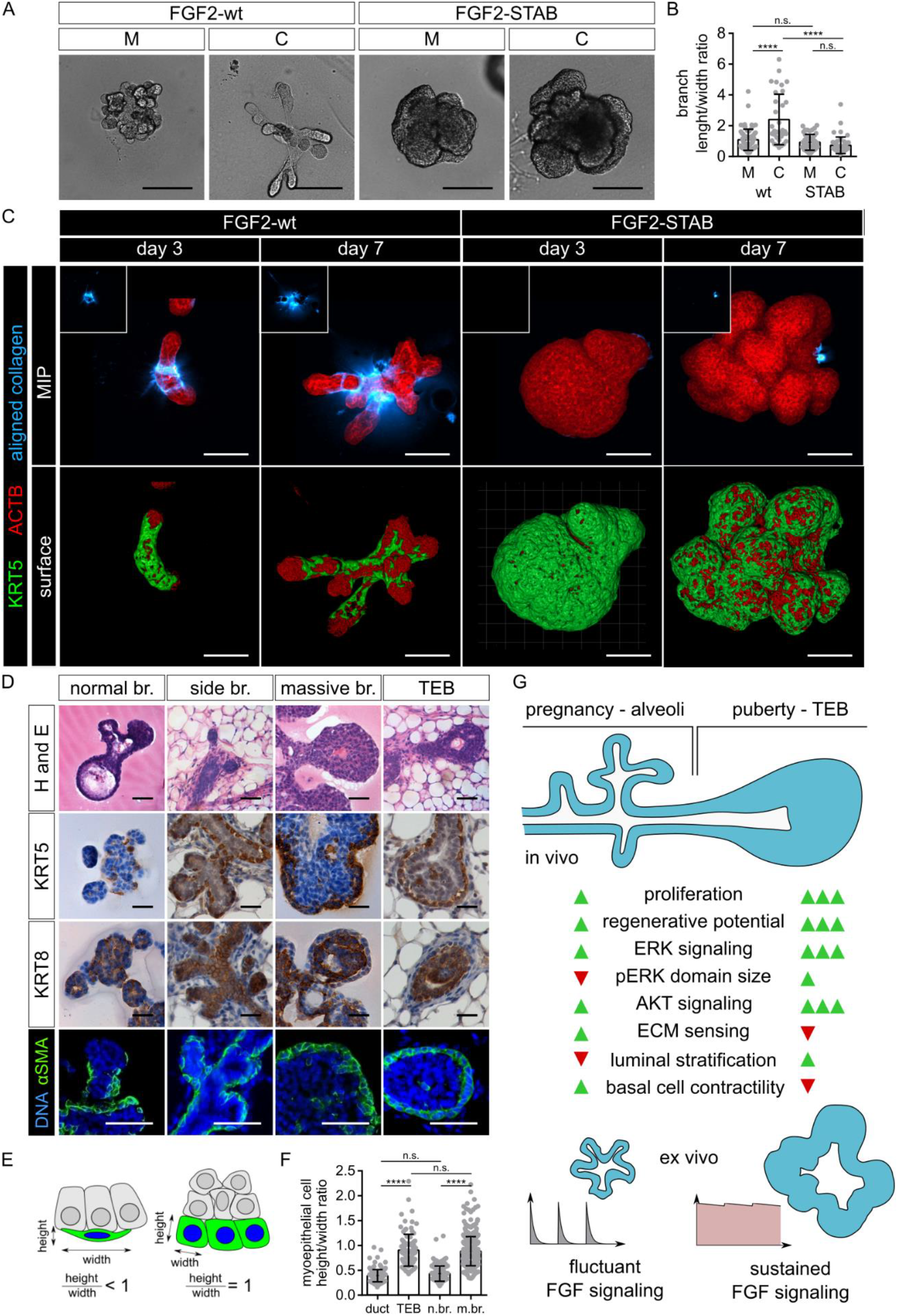
Massive branches phenocopy TEBs. **A**. Organoid morphogenesis in response to FGF2-wt and FGF2-STAB in Matrigel (M) or a mixture of Matrigel with collagen (C). Scale bars, 100 μm. Representative results of 3 independent biological replicates. **B**. Quantification of branch elongation in a mixture of Matrigel with collagen as branch length to width ratio. The plot shows mean + s.d., N = 71 (wt, M), 34 (wt, C), 48 (STAB, M) and 50 (STAB, C) branches. **C**. Maximum intensity projection (MIP) and surface reconstruction images of organoids cultured in a mixture of Matrigel with collagen (fluorescently labelled) and imaged using a confocal microscope. Scale bars, 100 μm. **D**. Hematoxylin and eosin (H and E; top row), immunohistochemical (second and third row), and immunofluorescence (bottom row) staining on sections of organoids with normal or massive branches, and of side branches and TEB in mammary gland tissue. Scale bars, 100 μm. **E**. Scheme depicting morphology of myoepithelial cells (green) with height/width ratio lower or equal to 1. Luminal cells are shown as grey. **F**. Quantification of myoepithelial cell height to width ratio of mammary gland side branches and TEBs, and normal branches of FGF2-wt-treated organoid and massive branches of FGF2-STAB-treated organoids. The plot shows mean + s.d., N = 155 (duct), 98 (TEB), 128 (normal br.) and 423 (massive br.) myoepithelial cells. ****P < 0.0001; (one-way ANOVA). **G**. Schematics of cell behaviors and underlying signaling activities in the mammary gland and in ex vivo mammary organoid cultures upon different FGF signaling dynamics. MG, mammary gland; TEB, terminal end bud.

Furthermore, by histological examination we noticed that by their bulb shape and extensive stratification, the massive branches of FGF2-STAB-treated organoids morphologically resembled terminal end buds (TEBs) of mammary gland in vivo (Figure 5D), the structures that drive mammary epithelial branching morphogenesis during puberty (Paine and Lewis, 2017). FGF2-wt-treated organoids contained branches formed mostly from a bi-layered epithelium and only some tips of the branches contained more than two layers of cells. Thereby, they resembled side branches in the mammary gland in vivo (Figure 5D). Immunohistological analysis of epithelial cell markers further accentuated the similarities between the normal and massive branches ex vivo and side branches and TEBs in vivo, respectively (Figure 5D). Moreover, the myoepithelial cells in massive branches of FGF2-STAB-treated organoids displayed a cuboidal shape similar to cap cells of TEBs, while the myoepithelial cells in the FGF2-wt-treated organoids and in the mammary ducts/side branches in vivo were flatter, based on their height to width ratio on organoid and tissue sections (Figure 5E, F). Collectively, the histology, lack of massive branch elongation in collagenous ECM, and increased regenerative potential of FGF2-STAB-treated organoids (Figure 2H; Supplemental Figure 7A-D) suggest that the massive branches of FGF2-STAB-treated organoids are ex vivo counterparts TEBs in vivo. Thus, our findings propose that distinct signaling dynamics encode different morphogenetic outcomes of physiological relevance (Figure 5G).

## Discussion

In this study, we investigated the role of FGF signaling dynamics in epithelial morphogenesis using the model of mammary epithelial organoids. FGF signaling is a well-established regulator of mammary gland branching morphogenesis (Ewald et al., 2008; Lu et al., 2008; Parsa et al., 2008; Pond et al., 2013; Sumbal and Koledova, 2019; Zhang et al., 2014). In vivo studies using genetic mouse models revealed roles of particular FGFRs in mammary gland morphogenesis (Lu et al., 2008; Parsa et al., 2008). Ex vivo 3D cultures of mammary epithelial organoids have elucidated the effects of individual FGF ligands on mammary epithelium (Ewald et al., 2008; Fata et al., 2007; Simian et al., 2001; Zhang et al., 2014). Thus, the role of FGF signaling in mammary epithelial morphogenesis has been well defined on the qualitative level; however, the quantitative aspects of FGF signaling, including the role of FGF signaling dynamics, have not been studied.

Using two variants of FGF2 with very different protein stabilities, and different medium changing strategies we exposed organoids to various dynamics of FGF2 availability. We discovered that fluctuant FGF2 signaling induces formation of thin branches, and that sustained FGF2 signaling leads to formation of massive, wide branches. In comparison to fluctuant FGF2 signaling, sustained FGF2 signaling led to increased activity of all FGFR downstream signaling pathways. In particular, ERK activity was increased and prolonged in FGF2-STAB-treated organoids in comparison to FGF2-wt-treated organoids. ERK activity dynamics has been suggested to play a key role in regulating mammary epithelial branching morphogenesis in response to different growth factors. In mammary organoid culture, TGFα induced sustained ERK activation and epithelial branching, and FGF7 induced only transient ERK activation and epithelial growth (Fata et al., 2007). Our study corroborates and extends these findings by demonstrating that different ERK activity dynamics, induced downstream of the same receptor, regulate mammary epithelial patterning and morphogenesis. Our study is, however, limited by endpoint-type analytical approaches to ERK activity quantification (immunodetection on fixed samples or Western blot and qPCR analysis at defined timepoints). Future studies using ERK activity biosensors (de la Cova et al., 2017; Komatsu et al., 2011) would be very helpful to elucidate ERK signaling dynamics during mammary epithelial morphogenesis in a much greater detail. Such studies could also help to define the role of ERK signaling domains in epithelial branch patterning and elongation. We found that domains of active ERK were larger in FGF2-STAB-treated organoids than in FGF2-wt-treated organoids and coincided with branch formation sites, suggesting that the size of ERK activity domains determines the diameter of nascent branches. Our experimental data are in agreement with computational simulations of multicellular morphogenesis using reaction-diffusion patterning, in which activator concentration and patterning determine morphological outcome of branching (Okuda et al., 2018). It remains to be determined, how ERK signaling domains become specified within the mammary epithelium.

Previous reports suggested that spatial enrichment of active ERK at the tips of the branches drives branch elongation (Huebner et al., 2016). In our model, active ERK was spatially enriched in cells in distal tips of the branches during both normal and massive branching. However, the branches efficiently elongated only in the FGF2-wt-treated organoids. The phenotype was even more prominent in ECM composed of Matrigel with collagen I that was previously demonstrated to promote formation of significantly longer branches than pure Matrigel (Nguyen-Ngoc and Ewald, 2013). This suggests that under sustained FGF2 signaling, epithelial stratification is dominant over epithelial cell intercalation into basal surface, the mechanism required for epithelial tube elongation (Neumann et al., 2018). This is probably due to changes in mechanosignaling in myoepithelial cells and/or changes in their mechanical properties, as suggested by changes in myoepithelial cell geometry and lack of force-mediated collagen I assembly under sustained FGF2 signaling.

ERK dynamics alone does not predict the cellular outcome. Rather, cell fate depends on a combination of downstream signaling activities induced by particular growth factor dynamics (Chen et al., 2012; Sampattavanich et al., 2018). On tissue level we detected contribution of AKT-mediated cell activities to epithelial branching, and we identified AKT signaling as a crucial regulator of epithelial stratification and massive branching phenotype. Furthermore, we pinpointed the important role of “basal” AKT signaling, provided by insulin present in the basal medium during ex vivo culture. This is consistent with essential role of insulin-like growth factor 1 in ductal morphogenesis and formation of TEBs (Ruan and Kleinberg, 1999), the naturally occurring stratified mammary epithelial structures that drive mammary branching morphogenesis during puberty. Furthermore, our data demonstrate that inputs from several RTKs are integrated on downstream signaling nodes to regulate tissue morphogenesis in concert.

Importantly, our work brings novel insights how intracellular signaling activities of ERK and AKT combinatory regulate distinct morphological and functional states of mammary epithelium. Increased and prolonged ERK and AKT signaling promotes formation of TEB-like structures, while moderate and transient ERK and AKT signaling induces formation of thin branches. In our model these distinct downstream signaling activities and morphogenetic outcomes were induced by different FGF2 signaling dynamics and are in concordance with the essential roles of FGF signaling in TEB formation during puberty and in side branching and alveologenesis during pregnancy (Lu et al., 2008; Parsa et al., 2008). The potential mechanisms for regulation of FGF signaling dynamics in vivo include differential production, retention and distribution of FGF ligands in the mammary stroma, differential expression of FGFR isoforms, or differential use of downstream feedback loops (Soady et al., 2017). Furthermore, in vivo the distinct ERK and AKT signaling activities most likely result from combinatorial effects of several growth factors and other signals, such as matrix metalloproteinases, present in the mammary stroma during postnatal mammary gland development, which collectively regulate mammary gland morphogenesis (Gjorevski and Nelson, 2011; Mori et al., 2013).

Our study also revealed that the distinct morphological states of mammary epithelium differentially engage ECM to further support epithelial morphogenesis. The non-stratified, side branch-like epithelial structures exert mechanical forces on surrounding collagen, which promotes branch elongation. However, the highly stratified TEB-like epithelial structures do not engage collagen in the surrounding matrix. These findings are in agreement with substantial collagen organization around mammary ducts and necks of TEBs, and lack of organized collagen around TEBs in vivo (Brownfield et al., 2013; Hinck and Silberstein, 2005; Lilla and Werb, 2010).

Nevertheless, we acknowledge that by their high proliferation, decreased apoptosis, increased stem cell properties, and multi-layered architecture with intact myoepithelial cell layer, the massively branched epithelial structures induced by sustained FGF2 signaling resemble not only TEBs, but also early, non-invasive stages of breast tumors, including hyperplasia, and their in vitro models induced by oncogenic RTK-Ras signaling (Muthuswamy et al., 2001; Welm et al., 2002). This testifies to the common critical cellular and signaling mechanisms used in both morphogenesis and cancerogenesis.

Precise regulation of complex cell behaviors on population level is essential for building functional tissues and organs. It is achieved by cell communication codes that we are only beginning to unravel. Understanding of the signaling codes in development is required for our understanding of aberrant signaling in developmental defects and disease, including cancer, and development of effective therapies. Moreover, it is an essential prerequisite for tissue engineering, stem cell therapy and regenerative medicine. Our study brings insights on how FGF signal availability regulates epithelial branched pattern formation. Future studies using multi-dimensional measurements of intracellular signaling activities on tissue scale with single-cell resolution shall help decipher the cell communication codes, including the relationship between signal processing and cell fate decision-making, during organ development and morphogenesis.

## Methods

### Mice

Nulliparous ICR females, 6-8 weeks old were used in this study. The animals were obtained from the Laboratory Animal Breeding and Experimental Facility of the Faculty of Medicine, Masaryk University. Experiments involving animals were approved in accordance with the Ministry of Agriculture of the Czech Republic, and the Expert Committee for Laboratory Animal Welfare at the Faculty of Medicine, Masaryk University.

### Organoid isolation and culture

Organoid isolation was performed as previously described (Koledova and Lu, 2017). Briefly, the mice were euthanized by cervical dislocation, the mammary glands were removed, mechanically disintegrated and partially digested in a solution of collagenase and trypsin [2 mg/ml collagenase (Sigma), 2 mg/ml trypsin (Thermo Fisher Scientific), 5 μg/ml insulin (Sigma), 50 μg/ml gentamicin (Sigma), 5% fetal bovine serum (FBS; Hyclone) in DMEM/F12 (Thermo Fisher Scientific)] for 30 min at 37°C with shaking at 100 rpm. Resulting tissue suspension was treated with 20 U/ml DNase I (Sigma) and exposed to five rounds of differential centrifugation at 450 × g for 10 s, which resulted in separation of epithelial (organoid) and stromal fractions. The organoids were resuspended in basal organoid medium [BOM; 1× ITS, 100 U/ml of penicillin, and 100 μg/ml of streptomycin in DMEM/F12 (all from Thermo Fisher Scientific)] and counted.

Subsequently, the organoids were mixed with ECM, either pure Matrigel or a mixture of Matrigel with collagen I. The mixture of Matrigel with collagen I was prepared as previously published (Koledova, 2017). Briefly, pre-assembled neutralized collagen I was prepared by combing 12.5 volumes of collagen type I (Corning) with 1 volume of 0.22 M NaOH, 5× collagen reconstitution buffer (5× MEM, 20 μg/ml NaHCO_3_, 0.1 M Hepes), and DMEM/F12 to the final concentration 2.58 mg/ml collagen and incubation of the neutralized collagen I for 1.5 h on ice. Then pre-assembled collagen was mixed with Matrigel at the ratio 7:3 and this mixture was immediately used to plate organoids. Fluorescently labelled collagen I was prepared according to a published protocol (Geraldo et al., 2013) using TAMRA (Sigma).

The organoids in ECM were plated in 50 μl domes at following densities: 250-300 organoids per dome for time-lapse and whole mount immunofluorescent analysis, 300-500 organoids per dome for histological and transcriptional analysis, 500-1,000 organoids per dome for Western blot analysis. After setting the ECM for 45-60 min at 37°C, the cultures were overlaid with BOM supplied with no FGF, or with FGF2-wt or FGF2-STAB (Enantis) according to the experiment. Unless stated otherwise, concentration of FGF2 was 1 nM. The cultures were incubated in a humidified atmosphere of 5% CO_2_ at 37°C on Olympus IX81 microscope equipped with Hamamatsu camera and CellR system for time-lapse imaging. The organoids were photographed every 60 min for 9 days with manual refocusing every day. For analysis of cell proliferation, 10 μM BrdU (Sigma) was added to the medium 3 h prior to organoid culture fixation.

For long term culture, organoids were cultured for 30 days with media changed every 3 days. Then Matrigel with organoids was disrupted with a 1 ml pipette and treated with trypsin-EDTA for 5 min, passed through a 25-gauge needle to obtain single cells. 30,000 cells were seeded in 50 μl Matrigel and treated with BOM with appropriate growth factors. The cells were cultured for 16 days with media changed every 3 days.

For inhibitor assays, the organoid cultures were treated with inhibitors in concentrations as indicated (Supplemental Table 1). Fresh medium with 1 nM FGF2 and/or inhibitors was changed every 3 days if not indicated otherwise.

### Organoid morphology analysis

Organoid branching was evaluated in ImageJ from time-lapse videos and branching was defined as formation of a new bud/branch from the organoid. Branches wider than 150 μm were considered as “massive branches”. 20 organoids per condition per experiment were analyzed, organoids that fused with another organoid or collapsed after attachment to the bottom of the well were excluded from the quantification.

### Organoid histology, immunohistochemistry and immunofluorescence on histological sections

Organoid cultures were washed 3 times with PBS and fixed for 30 min in 4% paraformaldehyde (Electron Microscopy Sciences). After washing with PBS, the cultures were embedded in 3% low gelling temperature agarose (Sigma). After solidification, the samples were dehydrated and embedded in paraffin. Sections (5 μm thick) were cut and dewaxed for hematoxylin and eosin staining or immunostaining.

For immunohistochemistry, antigens were retrieved in Citrate buffer, pH 6 or Tris-EDTA buffer, pH 9 (both Dako), endogenous peroxidase activity was blocked using 3% hydrogen peroxide and sections were blocked in PBS with 10% FBS (blocking buffer) for 1 h. Then, sections were incubated with primary antibody in blocking buffer for 2 h. After washing, sections were incubated with secondary antibody in blocking buffer for 30 min. Nuclei were counterstained with Mayer’s hematoxylin, sections were dehydrated and mounted in Pertex (Histolab Products). The samples were photographed using Leica DM5000 equipped with Leica DFC480 camera.

To perform immunofluorescence staining, antigens were retrieved in Citrate buffer, pH 6 or Tris-EDTA buffer, pH9 (both Dako), blocked in PBS with 10% FBS (blocking buffer) for 1 h and incubated with primary antibodies in blocking buffer overnight. After washing, sections were incubated with secondary antibodies in blocking buffer for 2 h. Nuclei were counterstained with DAPI for 10 min and slides were mount with Mowiol (Sigma). The samples were photographed using Leica DM5000 equipped with Leica DFC480 camera or Zeiss Axioimager 2. Antibodies and their concentrations used in this study are listed in Supplemental Table 2.

### Whole-mount organoid staining

For whole-mount imaging, organoids were 3D cultured in coverslip-bottom dishes (Ibidi). Organoid cultures were fixed with 4% paraformaldehyde for 30 min, permeabilized in 0.05% Triton X-100 in PBS for 1 h and blocked for 3 h with blocking buffer. Primary antibodies (Supplemental Table 2) diluted in blocking buffer were incubated with samples overnight at 4°C. After washing, samples were incubated with secondary antibodies (Supplemental Table 2) and 2 U/sample phalloidin-AlexaFluor488 (Thermo Fisher Scientific) in blocking solution for 2 h in darkness. Subsequently, samples were stained with 0.5 μg/ml DAPI (Merck) for 10 min and stored in PBS in 4°C in darkness until analyzed. The organoids were imaged using an LSM800 confocal microscope (Zeiss) and analyzed and exported using ZEN blue software (Zeiss).

### Mammary gland processing for histology

For histological analysis, 4^th^ mammary glands were removed from euthanized mice, spread on microscopy slide and fixed overnight in 4% paraformaldehyde. After washing in tap water, mammary glands were moved to histological cassettes and processed via standard procedure for paraffin embedding. Paraffin sections were cut (5 μm thick), dewaxed using xylene and rehydrated for hematoxylin and eosin staining or immunostaining.

### Tranmission electron microscopy

The samples were fixed with 3% glutaraldehyde in 100 mM sodium cacodylate buffer, pH 7.4 for 45 min, postfixed in 1% OsO_4_ for 50 min, and washed with cacodylate buffer. After embedding in 1% agar blocks, the samples were dehydrated in increasing ethanol series (50, 70, 96, and 100%), treated with 100% acetone, and embedded in Durcupan resin (Merck). Ultrathin sections were prepared using LKB 8802A Ultramicrotome, stained with uranyl acetate and Reynold’s lead citrate (Merck), and examined with FEI Morgagni 286(D) transmission electron microscope.

### Mammosphere assay

To test primary mammosphere formation efficiency, organoids which had been treated with BOM only, 1 nM FGF2-wt or 1 nM FGF2-STAB, were resuspended on day 4 of culture in 5 mM EDTA in PBS and shaken at 200 rpm on orbital shaker on ice for 1 h to dissolve the Matrigel. After washing with PBS, organoids were treated with HyQtase (Hyclone/GE Healthcare) for 10 min at 37°C and disintegrated by passing through a 24-gauge needle to acquire single cell suspension. Cells were then resuspended in mammosphere medium [1×B27 without vitamin A, 100 U/ml of penicillin, 100 μg/ml of streptomycin (all Thermo Fisher Scientific), 4 μg/ml heparin (Sigma), 20 ng/ml epidermal growth factor (Peprotech), 10 ng/ml FGF2-wt (Enantis) in phenol red-free DMEM/F12 (Thermo Fisher Scientific)] and seeded in polyHEMA-coated 6-well plates in concentration 20,000 cells per well. Fresh medium was provided every 3 days. After 9 days of culture, mammospheres were counted and mammosphere formation efficiency was calculated as number of formed mammospheres divided by number of cells seeded × 100. To assess secondary and tertiary mammosphere formation efficiency, primary or secondary mammospheres, respectively, were treated with HyQtase for 10 min at 37°C and disintegrated by passing through a 24-gauge needle to acquire single cell suspension. 20,000 cells per well were seeded in polyHEMA-coated 6-well plates in mammosphere medium and cultured and quantified similarly as primary mammospheres.

### Western blot

Organoid cultures were disintegrated by pipetting up and down in ice cold PBS with phosphatase inhibitors (10 mM β-glycerophosphate, 5 mM NaF, 1 mM Na_3_VO_4_), spun down and lysed in RIPA buffer (150 mM NaCl, 1.0% NP-40, 0.5% sodium deoxycholate, 0.1% SDS, 50 mM Tris, pH 8.0) supplied with proteinase and phosphatase inhibitors (10 mM β-glycerophosphate, 5 mM NaF, 1 mM Na3VO4, 1 mM dithiotreitol, 0.5 mM phenylmethanesulphonylfluoride, 2 μg/ml aprotinin, 10 μg/ml leupeptin; all Merck). Protein lysates were homogenized by sonication, cleared by centrifugation and protein concentration was measured using the Bradford reagent. Denatured, reduced samples were resolved on 10% SDS-PAGE gels and blotted onto PVDF membranes (Merck). Membranes were blocked with 5% non-fat milk in PBS with 0.05% Tween-20 (Merck; blocking buffer) and incubated with primary antibodies (Supplemental Table 2) diluted in blocking buffer overnight at 4°C. After washing in PBS with 0.05% Tween-20, membranes were incubated with horseradish peroxidase-conjugated secondary antibodies (anti-mouse antibody and anti-rabbit antibody, Cell Signaling Technology) for 1 h at room temperature. Signal was developed using an ECL substrate (100 mM Tris-HCl, pH 8.5, 0.2 mM coumaric acid, 1.25 mM luminol, 0.01% H_2_O_2_; all Merck) and exposed on X-ray films (Agfa), which were then scanned, and band density was analyzed using ImageJ. Phosphorylated and total proteins and actin were analyzed on a single blot.

### qRT-PCR

Organoid cultures were disintegrated by pippetting up and down in RLT buffer (Qiagen) and RNA was isolated using RNeasy Mini Kit (Qiagen) according to the manufacturer’s instruction. RNA concentration was measured using NanoDrop 2000 (Thermo Fisher Scientific). RNA was transcribed into cDNA by using Transcriptor First Strand cDNA Synthesis Kit (Roche) or TaqMan Reverse Transcription kit (Life Technologies). Real-time qPCR was performed using 5 ng cDNA, 5 pmol of the forward and reverse gene-specific primers each (Supplemental Table 3) in Light Cycler SYBR Green I Master mix (Roche) on LightCycler 480 II (Roche). Relative gene expression was calculated using the ∆∆Ct method and normalization to two housekeeping genes, β-actin (*Actb*) and Eukaryotic elongation factor 1 γ (*Eef1g*).

### Mathematical modeling of FGF2 concentration dynamics

Mathematical modeling of FGF2 concentration dynamics was based on half-lives of FGF2 variants: 6 h for FGF2-wt and c.a. 720 h for FGF2-STAB) (Dvorak et al., 2018). Function describing change in concentration (y) dependent on time (x) was determined as shown in (A1).

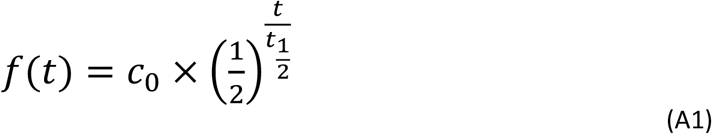

In (A1) c_0_ is initial concentration of FGF2, t is time in hours and t_1/2_ is half-life of FGF2. For conditions of 1 nM FGF with media not changed, function of FGF2 concentration dynamics was determined for FGF2-wt and FGF2-STAB, respectively, as shown in (A2).

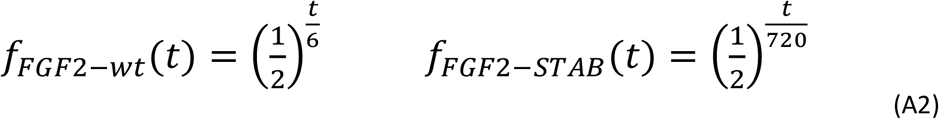

The area under the curve (AUC) was calculated as a sum of definite integrals with defined intervals as shown in (A3),

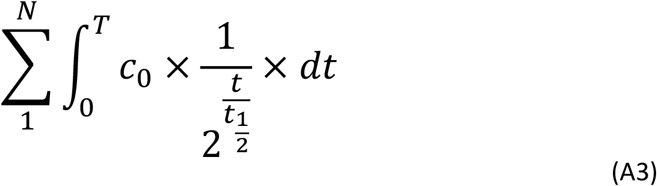

Where N is number of media changes (1 for NC; 3 for C3d; 9 for Ced; 36 for C6h); T is a time period determining the duration between two media changes (216 for NC; 72 for C3d; 24 for Ced; 6 for C6h); c_0_ is initial concentration; t is time and t_1/2_ is half-life of FGF2.

For experiments where medium with FGF2 was washed out, FGF concentration after the washout was determined as zero. For experiment where FGF2 ligands were added every 6 h without changing the medium, the AUC was calculated as a sum of definite integrals of single 6 h intervals with the c_0_=a determined for each interval separately as a function of a sequence given as shown in (A4) and (A5).

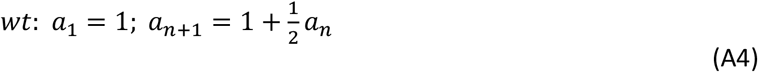

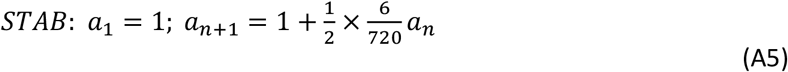

### Statistical analysis

Statistical analysis was performed using Prism software (GraphPad) using unpaired Student’s t-test or ANOVA. *P < 0.05, **P < 0.01, ***P < 0.001, ****P < 0.0001. Line plots and bar graphs were generated by Prism or Microsoft Excel and show mean ± standard deviation (s.d.).

## Acknowledgement

This research was supported by the Grant Agency of Masaryk University (grant no. MUNI/G/1446/2018 to Z.K and by funds from the Faculty of Medicine Masaryk University to junior researcher (Zuzana Koledova, ROZV/28/LF/2020). J.S. was funded by the P-Pool (Masaryk University, Faculty of Medicine) and by the Grant Agency of Masaryk University (grant no. MUNI/A/1565/2018). We thank Katarina Mareckova for excellent histology technical support, Dobromila Klemova for help with preparation of sections for electron microscopy, and Anas Rabata for help with mammosphere assay. We thank Veronika Stepankova, Radka Chaloupkova and Jiri Damborsky for providing the FGF2-wt and FGF2-STAB. We acknowledge the core facility CELLIM of CEITEC supported by the Czech-BioImaging large RI project (LM2018129 funded by MEYS CR) for their support with obtaining scientific data presented in this paper.

## Author Contributions

J.S. designed and performed the experiments, analyzed the data and wrote the manuscript. Z.K. conceptualized the study, secured funding, designed and performed the experiments, analyzed the data and wrote the manuscript. T.V. performed immunostainings and analyzed the data. All authors approved the final manuscript.

## Data availability

Data supporting the findings of this work are available within the paper and its Supplemental Information files. Other data are available from the corresponding author upon reasonable request.

