## Supplementary information for "FGF signaling dynamics regulates epithelial patterning and morphogenesis"

Sumbal et al.

#### Supplemental Information

Supplemental Figures: 12

Supplemental Tables: 5

Supplemental Videos: 5

#### Supplemental Figures

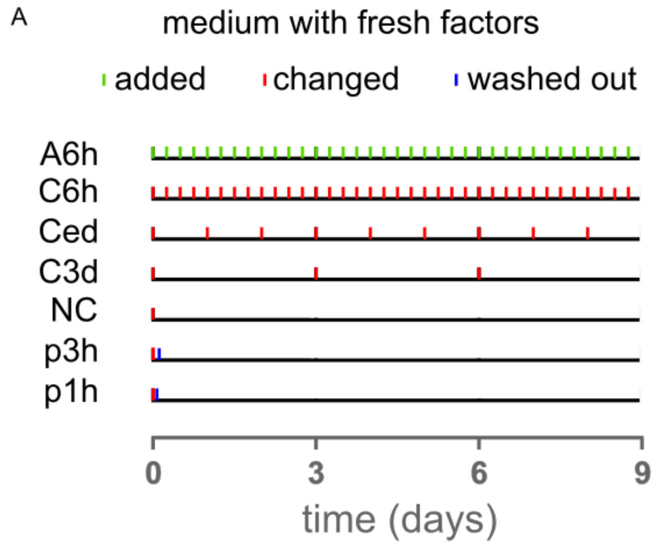


**Supplemental Figure 1. Schematic presentation of medium changing or growth factor adding strategies.**

**A.** P1h, 1-hour pulse; p3h, 3-hour pulse; NC, medium was not changed during the whole 9-day culture of organoids; C3d, medium was changed every 3 days; Ced, medium was changed every day; C6h, medium was changed every 6 h; A6h, fresh FGF2 was added to the medium every 6 h and every 3 days, the full medium volume was changed.

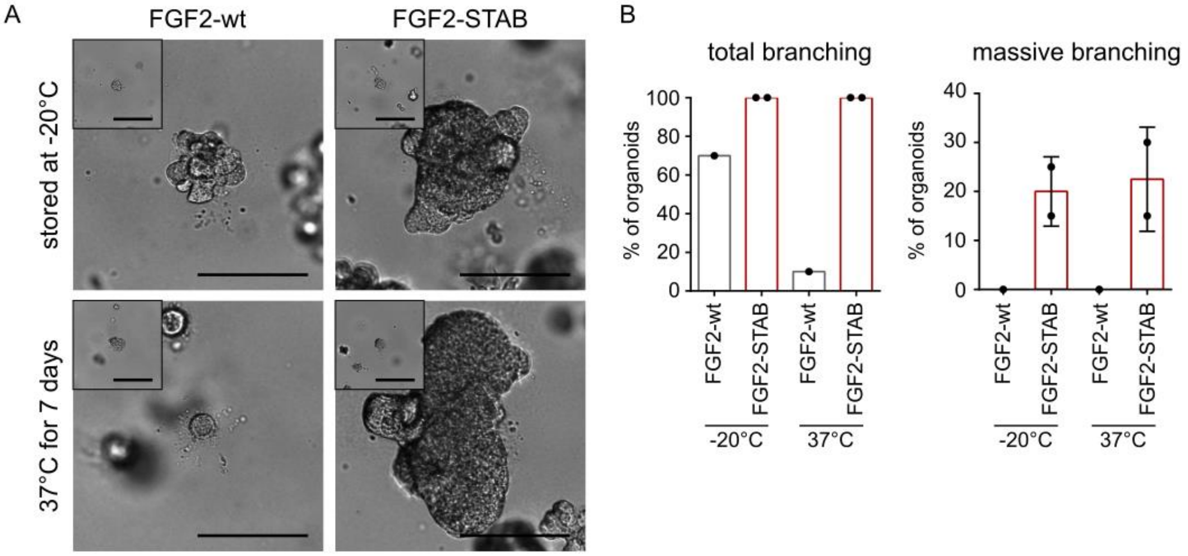


**Supplemental Figure 2**. **Thermostability of FGF2-STAB is demonstrated in mammary organoid culture.**

**A.** Images of organoids after 9 days of culture. FGF2-wt and FGF2-STAB were either stored at -20°C or preincubated at 37°C for 7 days before adding them in the culture medium. Insets show organoids on day 0 of culture. Scale bars, 200 µm.

**B.** Quantification of total branching and massive branching of organoids from **A**. The plots show mean + s.d., n = 1 (wt) and 2 (STAB) independent experiments, N = 20 organoids per experiment.

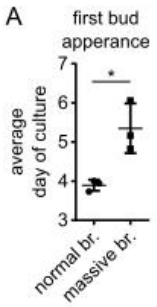


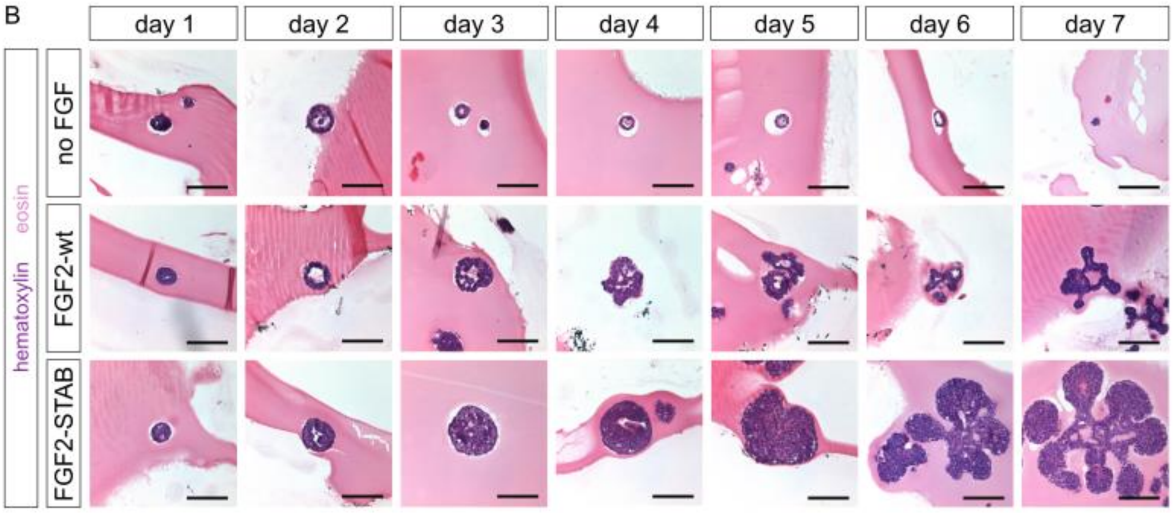


**Supplemental Figure 3**. **Branching is delayed in organoids exposed to sustained FGF signaling.**

**A.** Average day of the appearance of the first signs of branching was calculated from the experiment shown in Figure 2A. The plot shows mean + s.d., n = 3 independent experiments, N = 20 organoids per experiment. *P < 0.05 (Student's t-test).

A. **B.** Sections of organoids cultured with no FGF, with FGF2-wt or FGF2-STAB for 1 to 7 days stained with hematoxylin (blue) and eosin (pink). Scale bars, 200 µm. Details of FGF2-wt- and FGF2-STAB-treated organoids on day 7 of culture are shown in Figure 2C.

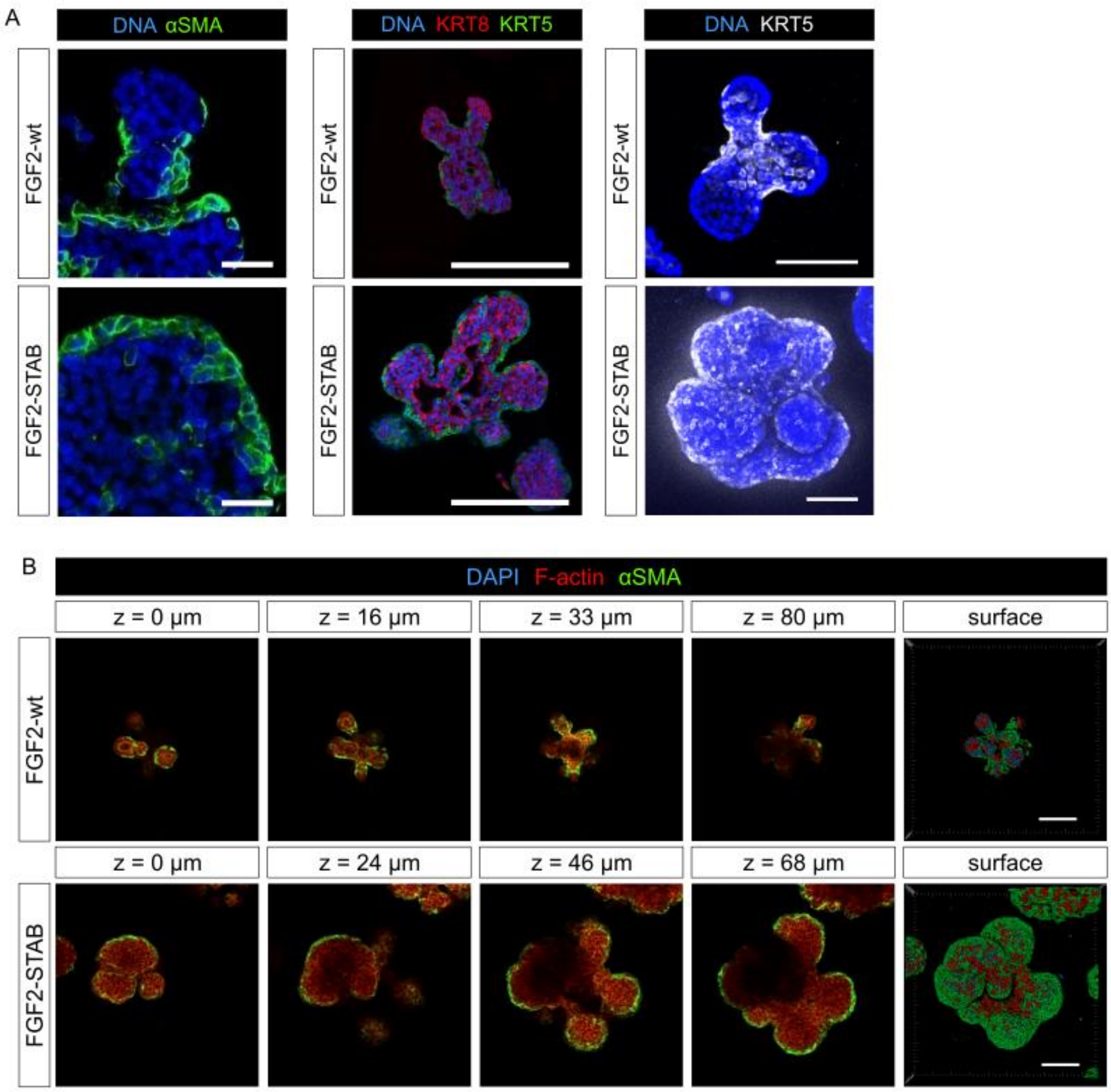


**Supplemental Figure 4**. **FGF2-STAB-treated organoids retain full myoepithelial coverage.**

**A.** Immunohistological (left, middle) and whole-mount (right) staining of FGF2-wt- or FGF2-STAB-treated organoids. αSMA, α smooth muscle actin; KRT5, keratin 5; KRT8, keratin 8. Scale bars, 100 µm.

**B.** Confocal stack frames and surface reconstruction images of organoids treated with FGF2-wt or FGF2-STAB for 6 days. DAPI, nuclei. Scale bars, 100 µm.

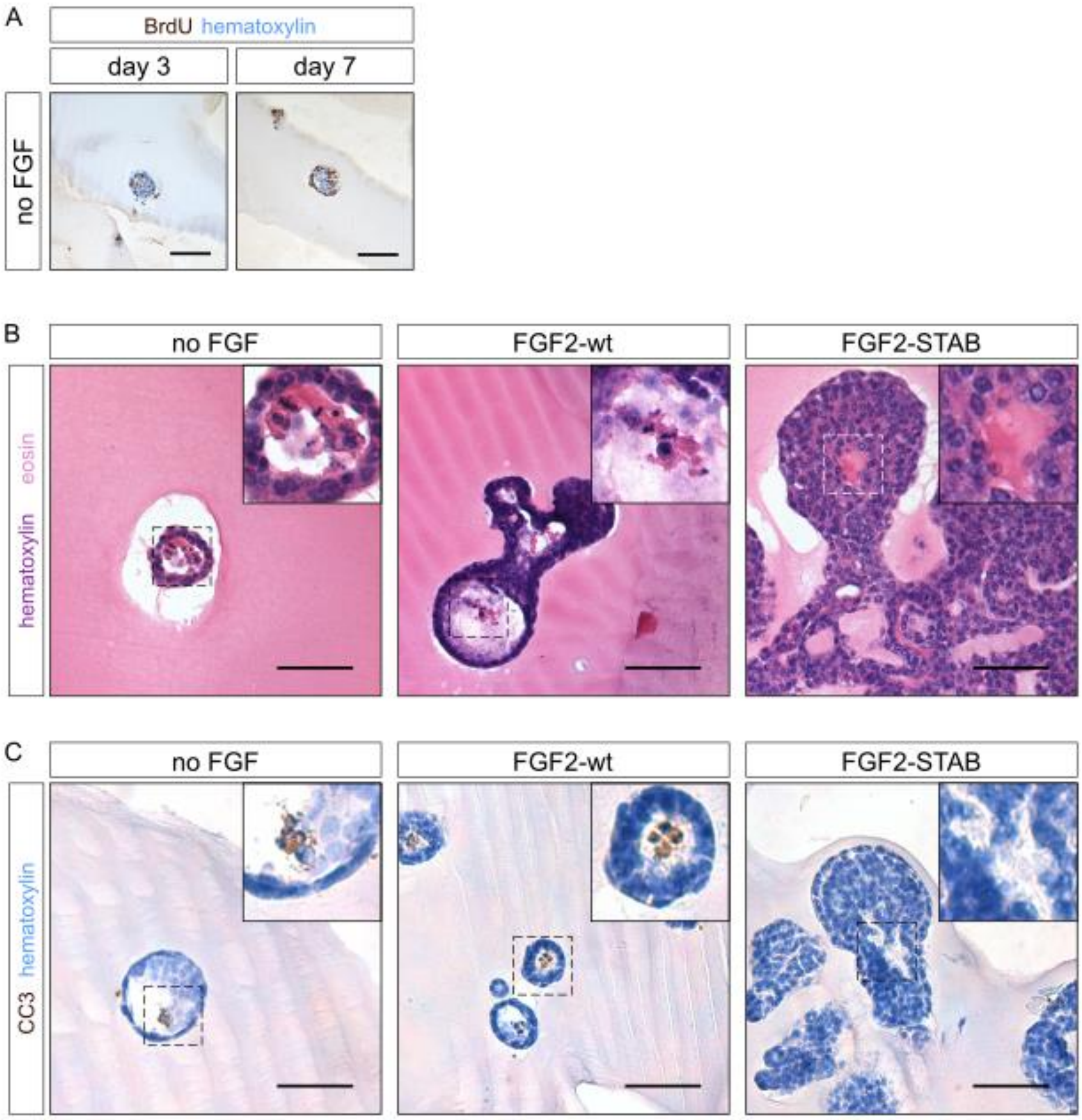


**Supplemental Figure 5. Sustained FGF2 signaling enhances cell survival.**

**A.** BrdU staining of control organoids to experiment in Figure 2F. Organoids were cultured without any FGF. Scale bars, 50 µm.

**B.** Sections of organoids treated with no FGF, with FGF2-wt or FGF2-STAB for 7 days stained with hematoxylin (blue) and eosin (pink). Apoptotic cells, shed to the lumen and with fractionated nuclei, are visible in organoids cultured with no FGF or with FGF2-wt. Scale bars, 100 µm.

**C.** Sections of organoids treated with no FGF, with FGF2-wt or FGF2-STAB for 7 days. CC3, cleaved caspase 3. Scale bars, 100 µm.

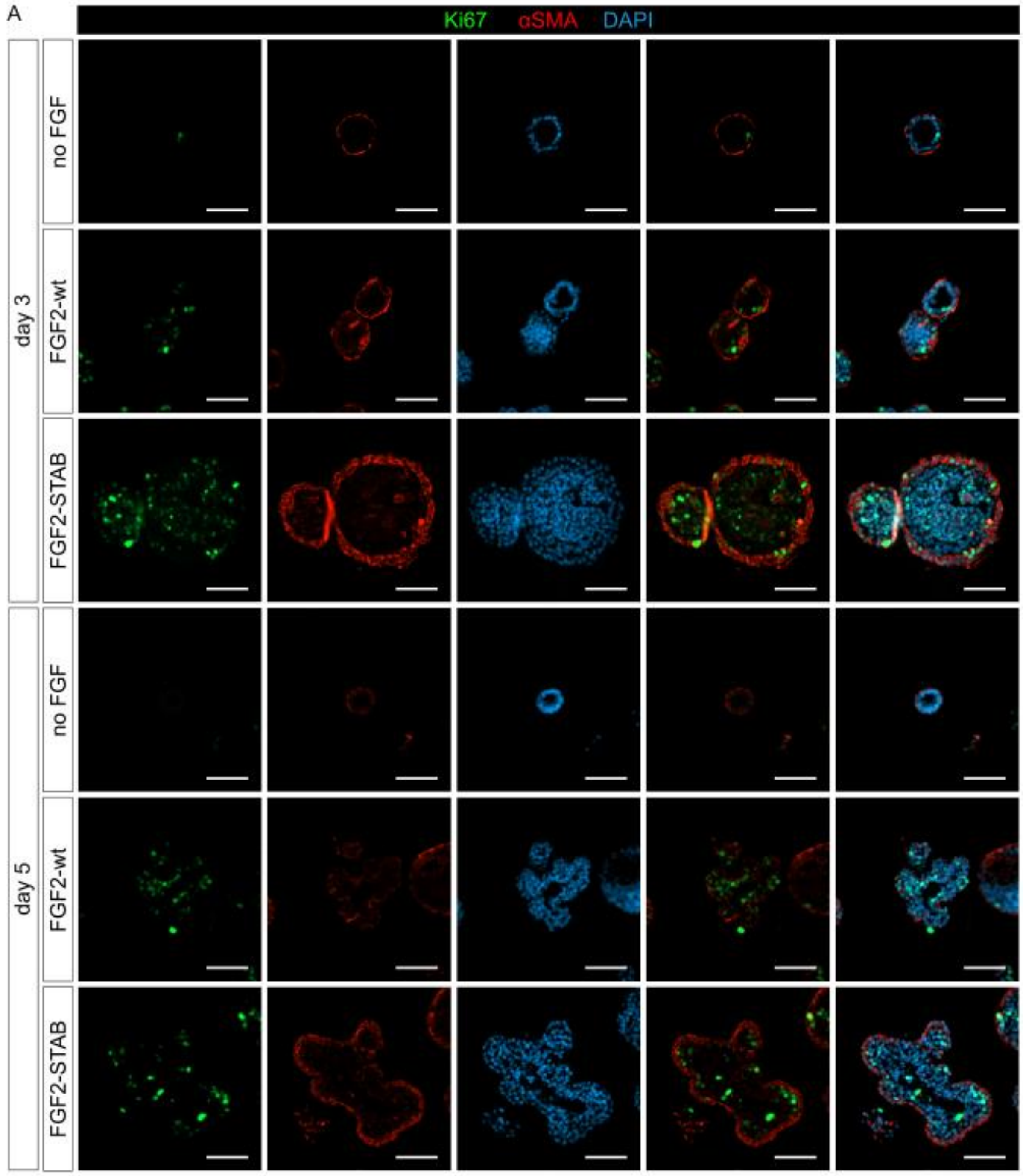


**Supplemental Figure 6. FGF2-STAB-treated organoids show enhanced proliferation.**

**A.** Extended data to Figure 2G. Immunohistological sections of organoids treated with no FGF, FGF2-wt or FGF2-STAB for 3 or 5 days. Scale bars, 50 µm.

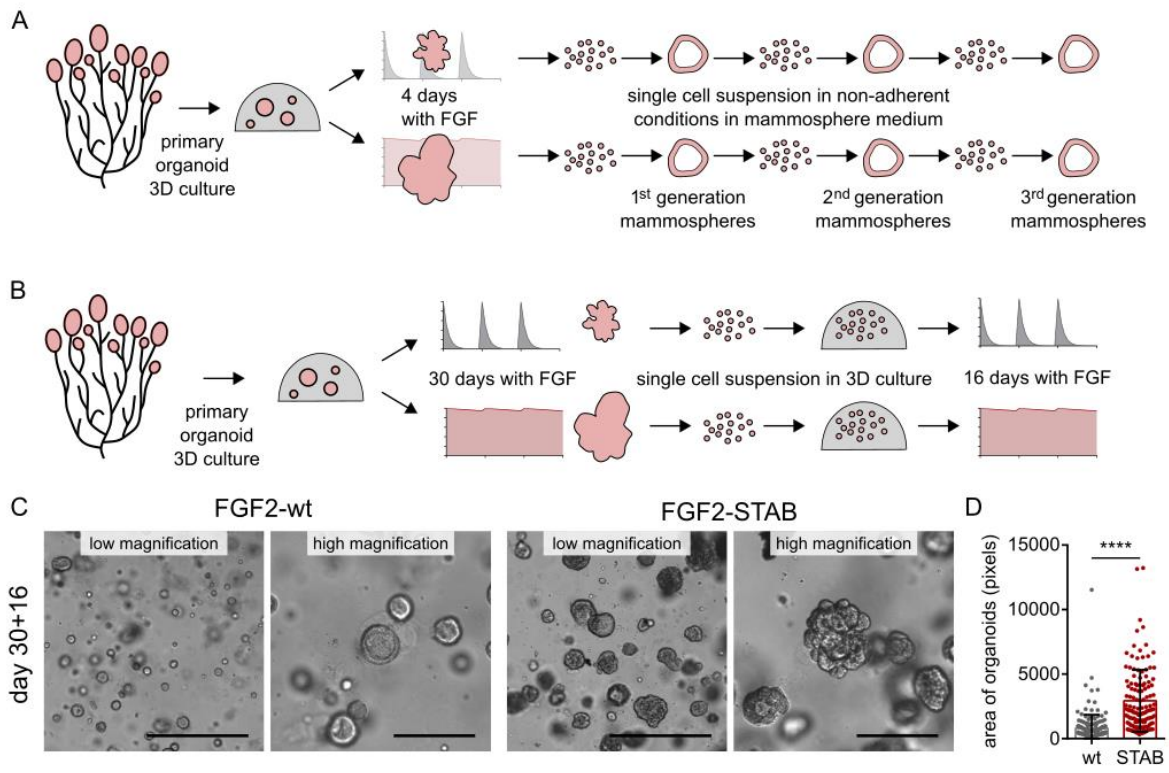


**Supplemental Figure 7. Sustained FGF signaling enhances regenerative capacity of organoids.**

**A.** Schematics of mammosphere formation experiment; results shown in Figure 2I. Primary organoids isolated from the mammary gland were subjected to either fluctuant or sustained FGF signaling, then dissociated to single cells and cultured in non-adherent condition for three generations.

**B.** Schematics of organoid formation assay. Primary organoids obtained from the mammary gland were subjected to either fluctuant or sustained FGF signaling, then dissociated to single cells, seeded in Matrigel and subjected to either fluctuant or sustained FGF signaling again. After 16 days, formation of organoids was assessed.

**C.** Representative appearance of 3D cultures of organoids obtained from single cells. Scale bars, 500 µm (low magnification) and 100 µm (high magnification).

**D.** Quantification of the size of organoids derived from single cells. The plot shows mean + s.d., n = 1 independent experiment, N = 186 (wt) and 129 (STAB) organoids. ****$P < 0.0001$ (Student's t-test).

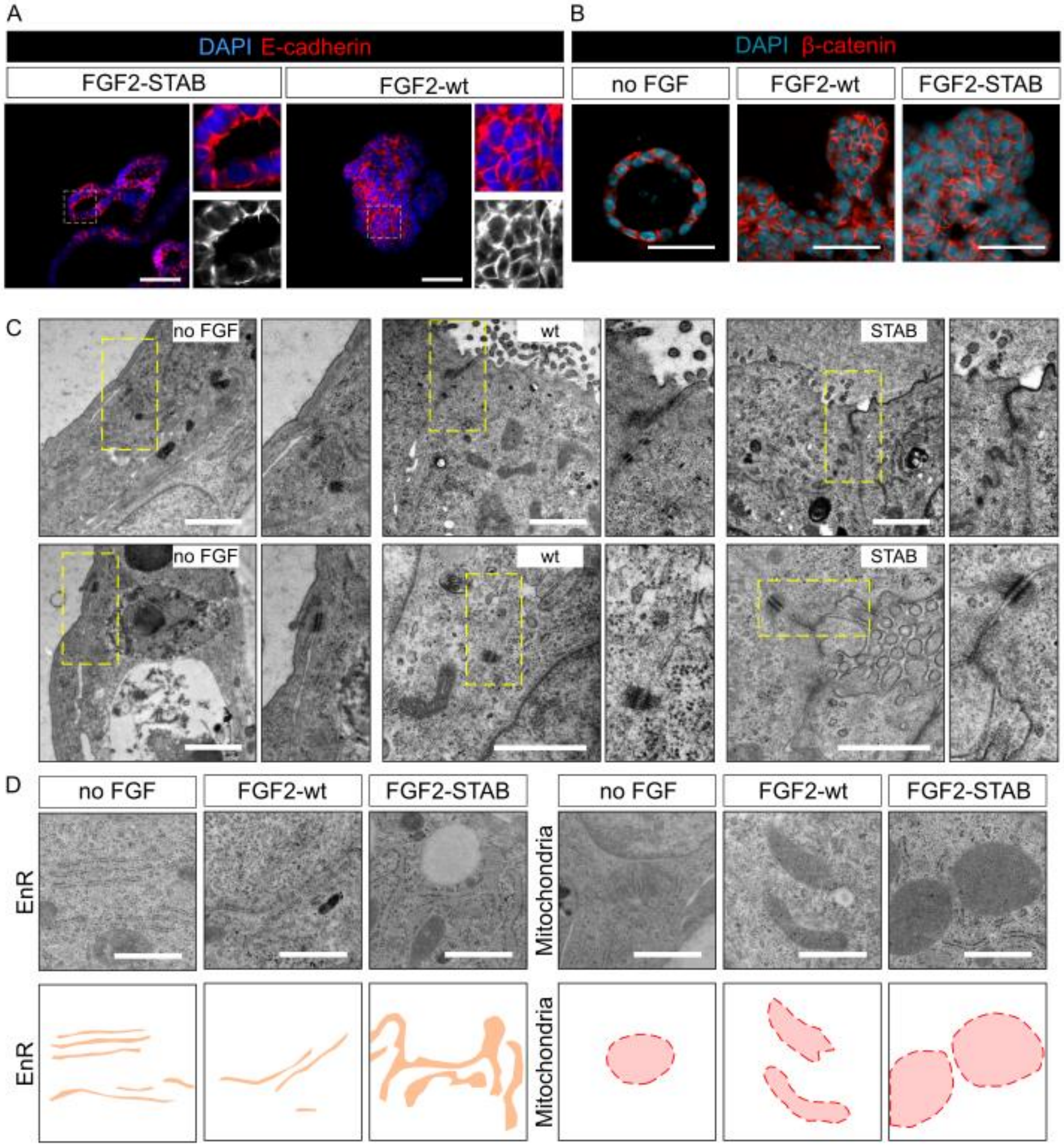


**Supplemental Figure 8. FGF2-STAB-induced epithelial stratification leads to decreased cell polarity.**

**A.** Immunohistology of organoids treated with FGF2-wt or FGF2-STAB for 7 days. Scale bars, 100 µm.

**B.** Immunohistology of organoids treated with no FGF, FGF2-wt or FGF2-STAB for 7 days. Scale bars, 100 µm.

**C.** TEM images of organoids treated with no FGF, with FGF2-wt or FGF2-STAB for 7 days. Upper panel shows the apical side and tight junctions of cells; lower panel shows desmosomes and microlumens. Marked area is shown in detail. Scale bars, 5 µm.

**D.** Ultrastructural analysis of cells in organoids treated with no FGF, with FGF2-wt or FGF2-STAB. TEM images (upper panel) and schematic presentation (lower panel) of endoplasmic reticulum (EnR) cisternae (orange) and mitochondria (red). Scale bar, 5 µm.

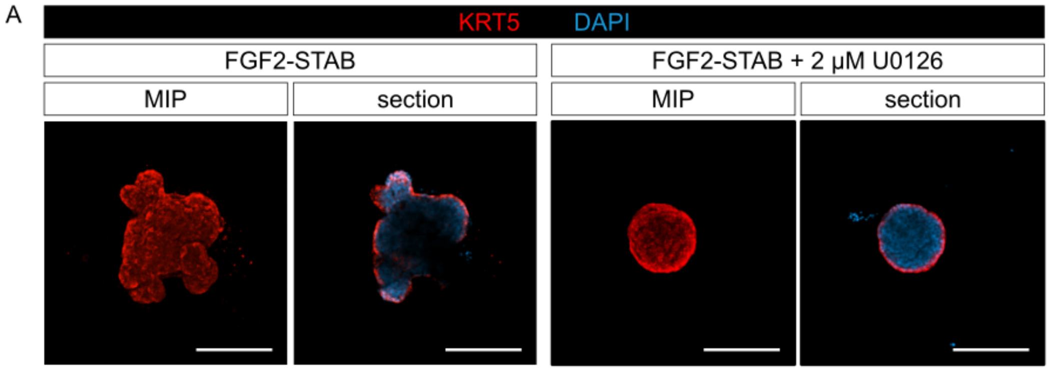


**Supplemental Figure 9. ERK pathway inhibition does not hinder full myoepithelial coverage of FGF2-STAB-treated organoids.**

**A.** Maximum intensity projection (MIP) and optical section images from confocal imaging of whole-mount organoids treated with FGF2-STAB and with no inhibitor or with 2 µM U0126 for 6 days. Scale bars, 200 µm.

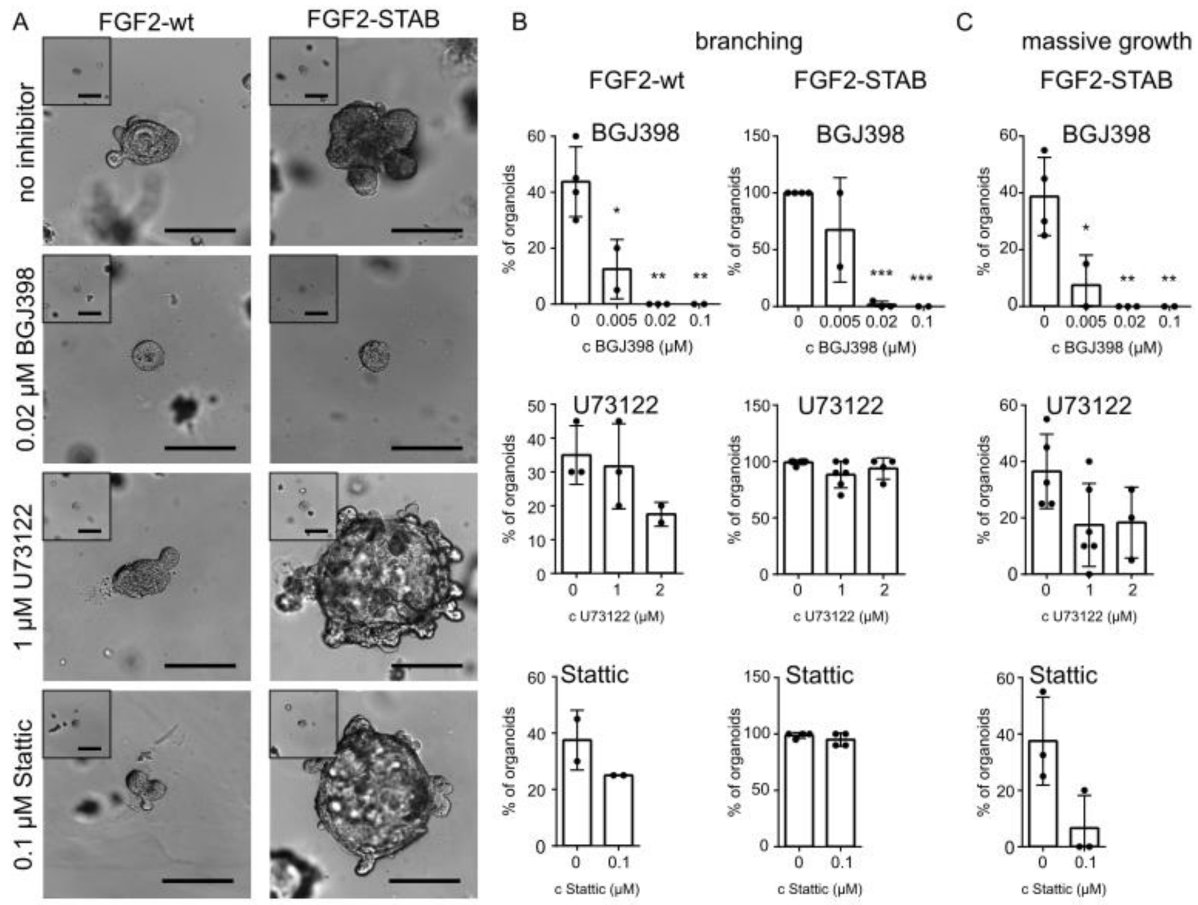


**Supplemental Figure 10. Inhibition of FGFR, PLCγ, or STAT3 interferes with FGF-induced morphogenesis.**

**A.** Representative appearance of organoids on day 9 of culture after being subjected to either fluctuant or sustained FGF signaling and treated with no inhibitor, 0.02 µM BGJ398, 1 µM U73122 or 0.1 µM Stattic. The images of organoids without inhibitor are the same as in Figure 3A because they are from the same experiment. Insets show organoids on day 0. Scale bars, 200 µm.

**B.** Quantification of total organoid branching after treatment with FGF2-wt or FGF2-STAB and with inhibitors at the indicated range. The plots show mean + s.d., n = 2-5 independent experiments, N =20 organoids per experiment. *P < 0.05; **P < 0.01; ***P < 0.001 (one-way ANOVA).

**C.** Quantification of massive growth of organoids after treatment with FGF2-STAB and with inhibitors at the indicated range. The plots show mean + s.d., n = 2-5 independent experiments, N =20 organoids per experiment. *P < 0.05; **P < 0.01 (one-way ANOVA).

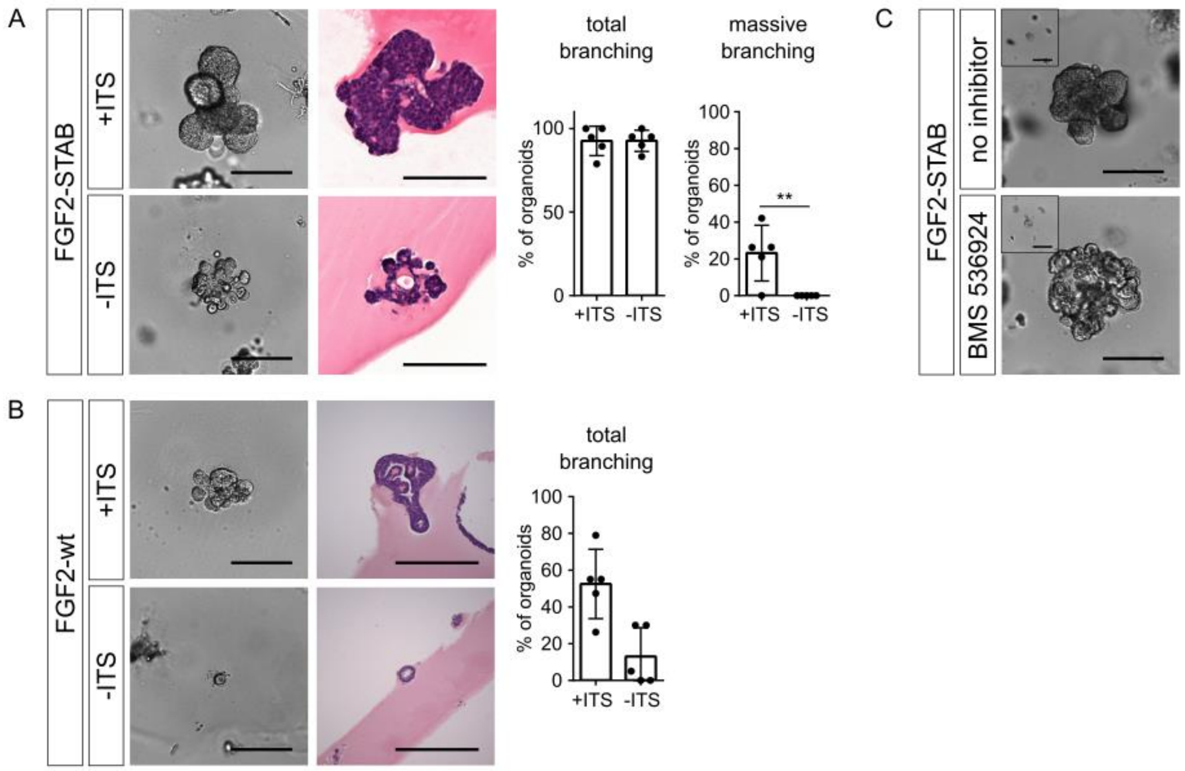


**Supplemental Figure 11. Insulin signaling supports FGF2-STAB-induced stratification.**

**A.** Organoid morphogenesis in response to FGF2-STAB in the presence or absence of insulin-transferrin-selenium (ITS) in basal organoid medium. Images of organoids after 9 days of culture, hematoxylin and eosin stained sections from the organoids, and quantification of total and massive branching. Scale bars, 200 µm. The plots show mean + s.d., n = 5 independent experiments, N = 20 organoids per experiment. **P < 0.01 (Student's t-test).

**B.** Organoid morphogenesis in response to FGF2-wt in the presence or absence of ITS in basal organoid medium. Images of organoids after 9 days of culture, hematoxylin and eosin stained sections from the organoids, and quantification of total branching. Scale bars, 200 µm. The plot shows mean + s.d., n = 5 independent experiments, N = 20 organoids per experiment. **P < 0.01 (Student's t-test). Scale bars, 200 µm.

**C.** Organoid morphogenesis in response to FGF2-STAB in the presence or absence of 0.1 µM BMS 536924 (insulin/IGF receptor inhibitor). Scale bars, 200 µm.

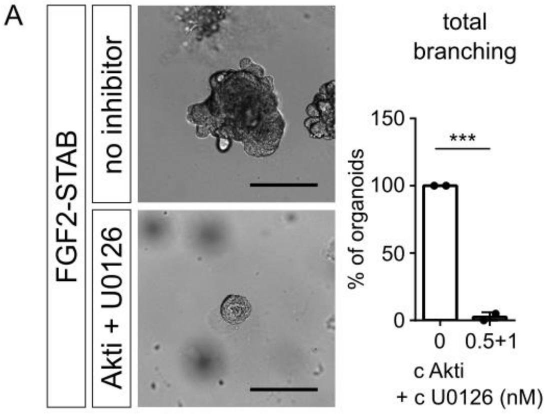


**Supplemental Figure 12. FGF signaling dynamics regulates epithelial morphogenesis through AKT and ERK signaling.**

**A.** Organoids cultured with FGF2-STAB in the presence or absence of 0.5 µM Akti1/2 and 1 µM U0126 for 9 days. Scale bars, 200 µm. The plots show quantification of total branching as mean + s.d., n = 2 independent experiments, N = 20 organoids per experiment. ***P < 0.001 (unpaired Student's t-test).

### Supplemental Tables

**Supplemental table 1. A list of small molecule inhibitors used.**

| Small molecule inhibitors | | | | | |
|---|---|---|---|---|---|
| **Inhibitor** | **Target** | **Source** | **Identifier** | **CAS** | **Dilution** |
| Akti1/2 | AKT1/2 | Tocris | 5773 | 612847-09-3 | 0.5 µM; 1 µM; 2 µM |
| BGJ398 | FGFR | SelleckChem | S2183 | 872511-34-7 | 0.1 µM; 0.02 µM; 0.005 µM |
| BMS536924 | IR/IGFR | Tocris | 4774 | 468740-43-4 | 0.1 µM |
| Stattic | STAT3 | Tocris | 2798 | 19983-44-9 | 0.1 µM |
| U0126 | MEK | Focus Biomolecules | 10-2179 | 109511-58-2 | 0.5 µM; 1 µM; 2 µM |
| Y73122 | PLCγ | Focus Biomolecules | 10-1061-0005 | 112648-68-7 | 1 µM; 2 µM |

**Supplemental table 2. A list of antibodies used in the study.**

| Primary Antibodies | | | |
|---|---|---|---|
| **Antibody** | **Source** | **Identifier** | **Dilution** |
| mouse monoclonal anti β-actin (C4) | Santa Cruz Biotechnology | sc-47778 | WB 1:1000 |
| rabbit polyclonal anti Akt | Cell Signalling Technology | #9272 | WB 1:1000 |
| rabbit polyclonal anti phospho-Akt1/2/3 (Ser473) | Santa Cruz Biotechnology | sc-7685-R | WB 1:1000 |
| mouse monoclonal anti BrdU (BU-33) | Sigma | B2531 | IHC-P 1:1000 |
| mouse monoclonal anti β-catenin (12F7) | Santa Cruz Biotechnology | sc-59737 | IHC-P IF 1:100 |
| rabbit polyclonal anti cleaved caspase 3 (Asp175) | Cell Signalling Technology | #9661 | IHC-P 1:200 |
| rat monoclonal anti E-cadherin | Santa Cruz Biotechnology | sc-59778 | IHC-P IF 1:100 |
| rabbit polyclonal anti ERK1/2 | Cell SignA13alling Technology | #9102 | WB 1:1000 |
| rabbit polyclonal anti keratin 5 | BioLegend | #905504 | IHC-P, IHC-P IF 1:250; IF 1:250 |
| mouse polyclonal anti keratin 8 | BioLegend | #904804 | IHC-P, IHC-P IF 1:250 |
| rabbit monoclonal anti Ki67 | Zytomed | RBK027-05 | IHC-P IF 1:500 |
| rabbit polyclonal anti phospho ERK1/2 (Thr202/Tyr 204) | Santa Cruz Biotechnology | sc-16982 | IHC-P, IF 1:250; WB 1:1000 |
| mouse monoclonal anti α Smooth muscle actin | Sigma | A2547 | IHC-P IF 1:200; IF 1:250 |

| Secondary Antibodies | | | |
|---|---|---|---|
| **Antibody** | **Source** | **Identifier** | **Dilution** |
| Goat anti-Mouse IgG (H+L), AlexaFluor 488 | Thermo Fisher Scientific | A11001 | 1:800 |
| Goat anti-Mouse IgG (H+L), AlexaFluor 568 | Thermo Fisher Scientific | A11031 | 1:800 |
| Goat anti-Rabbit IgG (H+L), AlexaFluor 488 | Thermo Fisher Scientific | A11008 | 1:800 |
| Goat anti-Rabbit IgG (H+L), AlexaFluor 568 | Thermo Fisher Scientific | A11011 | 1:800 |
| Goat anti-Rat IgG (H+L), AlexaFluor 555 | Thermo Fisher Scientific | A-21434 | 1:800 |
| EnVision+ Dual Link System-HRP | Dako | K4063 | N/A |
| anti-rabbit IgG, HRP-linked | Cell Signalling Technology | #7074 | WB 1:1000 |

**Supplemental table 3. A list of primers used for qPCR.**

| Primers | | | |
|---|---|---|---|
| **Target gene** | **Forward (5'-3')** | **Reverse (3'-5')** | **length (bp)** |
| *Actb* | GGCTGTATTCCCCTCCATCG | CCAGTTGGTAACAATGCCATGT | 154 |
| *Dusp6* | TCGGGCTGCTGCTCAAGAAAC | CGGTCAAGGTCAGACTCAATGTCC | 214 |
| *Eef1g* | TTCCTGCCGGCAAGGTTCCA | TGCCGCCTCTGGCGTACTTC | 119 |
| *Etv4* | CCACCAGGATCAAGAAGGAA | TTGTCTGGGGGAGTCATAGG | 131 |
| *Etv5* | AGGACCCCAGGCTGTACTTT | TGGCCGATTCTTCTGGATAC | 260 |

**Supplemental table 4. Numbers of experimental replicates for organoid branching assays.**

| number of replicates | | | | | |
|---|---|---|---|---|---|
| inhibitor | concentration | wt | | STAB | |
| | | experiments | organoids | experiments | organoids |
| U0126 | 0 | 4 | 80 | 4 | 80 |
| | 0.5 | 2 | 40 | 2 | 40 |
| | 1 | 3 | 60 | 3 | 60 |
| | 2 | 3 | 60 | 2 | 40 |
| Akti1/2 | 0 | 2 | 40 | 4 | 80 |
| | 0.5 | 2 | 40 | 3 | 60 |
| | 1 | 2 | 40 | 5 | 100 |
| | 2 | 2 | 40 | 5 | 100 |
| BGJ398 | 0 | 4 | 80 | 4 | 80 |
| | 0.005 | 2 | 40 | 4 | 80 |
| | 0.02 | 3 | 60 | 3 | 60 |
| | 0.1 | 2 | 40 | 3 | 60 |
| U73122 | 0 | 3 | 60 | 6 | 120 |
| | 1 | 3 | 60 | 6 | 120 |
| | 2 | 2 | 40 | 4 | 80 |
| Stattic | 0 | 2 | 40 | 4 | 80 |
| | 0.1 | 2 | 40 | 4 | 80 |
| FGF concentrations (nM) | | experiments | organoids | experiments | organoids |
| 0 | | 8 | 160 | 8 | 160 |
| 0.01 | | 2 | 40 | 2 | 40 |
| 0.025 | | 2 | 40 | 2 | 40 |
| 0.1 | | 3 | 60 | 3 | 60 |
| 0.25 | | 3 | 60 | 3 | 60 |
| 1 | | 8 | 160 | 9 | 180 |
| 5 | | 3 | 60 | 3 | 60 |
| 20 | | 2 | 40 | 1 | 20 |
| 40 | | 1 | 20 | 1 | 20 |
| Media changing | | experiments | organoids | experiments | organoids |
| p1h | | 1 | 20 | 2 | 40 |
| p3h | | 1 | 20 | 2 | 40 |
| NC | | 2 | 40 | 2 | 40 |
| C3d | | 5 | 100 | 4 | 80 |
| Ced | | 1 | 20 | 1 | 20 |
| C6h | | 3 | 60 | 4 | 80 |
| A6h | | 2 | 40 | 2 | 40 |

**Supplemental table 5. Numbers of tissue section analyzed in Figure 2E.**

| histology - nuclei counting | | | |
|---|---|---|---|
| | no FGF | FGF2-wt | FGF2-STAB |
| day 1 | 3 | 9 | 20 |
| day 2 | 5 | 11 | 20 |
| day 3 | 5 | 30 | 22 |
| day 4 | 24 | 36 | 20 |
| day 5 | 18 | 26 | 23 |
| day 6 | 20 | 25 | 34 |
| day 7 | 8 | 26 | 30 |
| day 8 | 13 | 34 | 40 |
| day 9 | 12 | 20 | 22 |
| **histology - Ki67** | | | |
| | no FGF | FGF2-wt | FGF2-STAB |
| day 3 | 13 | 14 | 25 |
| day 5 | 11 | 17 | 50 |

#### Supplemental Videos

**Supplemental Video 1.** Morphogenesis of mammary organoid over 9 days in Matrigel, cultured with basal organoid medium with no FGF. The medium was changed every 3 days.

**Supplemental Video 2.** Morphogenesis of mammary organoid over 9 days in Matrigel, cultured with basal organoid medium with 1 nM FGF2-wt. The medium was changed every 3 days.

**Supplemental Video 3.** Morphogenesis of mammary organoid over 9 days in Matrigel, cultured with basal organoid medium with 1 nM FGF2-STAB. The medium was changed every 3 days.

**Supplemental Video 4.** Morphogenesis of mammary organoid over 9 days in a mixture of Matrigel with collagen, cultured with basal organoid medium with 1 nM FGF2-wt. The medium was changed every 3 days.

**Supplemental Video 5.** Morphogenesis of mammary organoid over 9 days in a mixture of Matrigel with collagen, cultured with basal organoid medium with 1 nM FGF2-STAB. The medium was changed every 3 days.
